# Survival of a threatened salmon is linked to spatial variability in river conditions

**DOI:** 10.1101/2021.08.24.456882

**Authors:** Colby L. Hause, Gabriel P. Singer, Rebecca A. Buchanan, Dennis E. Cocherell, Nann A. Fangue, Andrew L. Rypel

**Author notes:** Corresponding Author: Colby Hause.

## Abstract

Extirpation of the Central Valley spring-run Chinook Salmon ESU (*Oncorhynchus tshawytscha*) from the San Joaquin River is emblematic of salmonid declines across the Pacific Northwest. Habitat restoration and fish reintroduction efforts are ongoing, but recent telemetry studies have revealed low outmigration survival of juveniles to the ocean. Previous investigations have focused on modeling survival relative to river discharge and geographic regions, but have largely overlooked the effects of habitat variability. To evaluate the link between environmental conditions and survival of juvenile spring-run Chinook Salmon, we combined high spatial resolution habitat mapping approaches with acoustic telemetry along a 150 km section of the San Joaquin River during the spring of 2019. While overall outmigration survival was low (5%), our habitat-based classification scheme described variation in survival of acoustic-tagged smolts better than other candidate models based on geography or distance. There were two regional mortality sinks evident along the longitudinal profile of the river, revealing poor survival in areas that shared warmer temperatures but that diverged in chlorophyll-*α*, fDOM, turbidity and dissolved oxygen levels. These findings demonstrate the value of integrating river habitat classification frameworks to improve our understanding of survival dynamics of imperiled fish populations. Importantly, our data generation and modeling methods can be applied to a wide variety of fish species that transit heterogeneous and diverse habitat types.

## Introduction

Pacific salmon populations native to California, USA, have declined throughout the last century, shifting once economically-viable runs to critically low numbers (Katz et al. 2013; Yoshiyama et al. 1998). Of the four currently recognized salmonid evolutionary significant units (ESUs) endemic to California’s Central Valley, three are listed under the Endangered Species Act (ESA), including spring-run Chinook Salmon *Oncorhynchus tshawytscha* (Moyle et al. 2017). Population declines in the Central Valley were initially driven by overharvest but have been exacerbated by systematic degradation of freshwater and estuarine habitats (Yoshiyama et al. 2000; Lund et al. 2007; Fisher 1994). Sexually-immature spring-run Chinook Salmon adults rely on early migration (January-February) during wet conditions to reach high-elevation tributaries that are otherwise inaccessible to salmon throughout most of the year (Fry 1961; Moyle et al. 2017). Fragmentation of rivers by large dams has been particularly devastating for this ESU, as these structures preclude fish from accessing critical cold-water summer holding areas where they mature before spawning in the fall. After emergence, juveniles migrate to the ocean during late spring or remain in freshwater for an additional year and exit as yearlings (Moyle 2002). Loss of essential habitats to meet life history requirements of the Central Valley spring-run ESU has resulted in their extirpation from the San Joaquin River (Yoshiyama et al. 2001). Recent efforts to reintroduce an experimental population back to this system (*Natural Resource Defense Council v. USBOR,* 2006; SJRRP 2018) revealed low juvenile survival to the ocean (Singer et al. in prep^1^). These results are consistent with previous studies estimating survival of acoustic-tagged juvenile fall-run Chinook Salmon in the same system (Buchanan et al. 2013, 2018).

Improved understanding of the habitats that promote or impair early life history success and overall cohort health is considered critical to the sustainable management of anadromous fish populations (Sass et al. 2017, Henderson et al. 2019, Michel 2019). Outmigration survival of juvenile salmon is tightly coupled to landscape characteristics as smolts undergo physiological changes while navigating large distances from riverine spawning grounds to ocean entry (Nislow and Armstrong 2012; Fausch et al. 2002). Furthermore, heterogeneity in habitat quality is thought to play an important role in shaping juvenile salmon ecology (e.g. feeding, rearing, and migration) (Frissell et al. 1986; Steel et al. 2012). Of the many physical and biological features that serve as measures of habitat quality, water chemistry is thought to have large impacts on salmon population dynamics (Zabel and Achord 2004; Warren et al. 1973; Clark et al. 1981). Nevertheless, previous efforts to improve outmigration success in the Central Valley have largely ignored habitat and instead focused on identifying the dominant physical environmental factors and geographic patterns that shape survival (Buchanan et al. 2013, 2018; Buchanan and Skalski 2020; Perry et al. 2010, 2018; Singer et al. in prep^1^). This approach has resulted in an incomplete understanding of how salmon interact with altered habitats across various spatial and temporal scales.

Several challenges have prevented researchers from incorporating water chemistry covariates or other measures of habitat quality into models of salmon migration survival. Spatially-detailed hydrologic and limnological data are difficult to collect and require more intense sampling than feasible by networks of stationary monitors. Additionally, fine-scale complexity of freshwater habitats that span large spatial extents presents challenges in accurately characterizing these landscapes (Fausch et al. 2002). For juvenile salmon migrating through a broad range of environmental conditions, gathering and characterizing information at multiple scales may improve our ability to relate key population metrics such as outmigration survival to habitat gradients.

The primary goals of this study were to (1) estimate survival of acoustic-tagged spring-run Chinook Salmon smolts from their release in the San Joaquin River through the Sacramento-San Joaquin River Delta (hereafter, “Delta”) to the Pacific Ocean, and (2) evaluate the link between landscape-level habitat variability and survival of smolts in the San Joaquin River. We employed a spatially-detailed habitat mapping method: fast limnological automated measurements (“FLAMe”, Crawford et al. 2015) to collect water chemistry data along a continuous transect of the mainstem San Joaquin River during the spring of 2019. We used FLAMe data in a novel approach to generate a river zonation framework for use with survival data from the concurrent acoustic telemetry study.

## Methods

### Study Area

We deployed a large array of Juvenile Salmon Acoustic Telemetry System (JSATS) receivers to track salmon movements from the lower San Joaquin River to ocean entry (27 sites, n = 133 receivers, Fig. 1). The study area for the telemetry array spanned 270 km of the San Joaquin River from the upper release location at Freemont Ford State Recreation Area (Merced County, CA, USA) to the Pacific Ocean at the Golden Gate Bridge (A17) (Fig. 1, Table 1). This region included the Delta, an inland tidal estuary largely formed by the Sacramento River to the north and the San Joaquin River to the south, ending at the confluence of these two rivers near Chipps Island (A15, Fig.1). The study area in the southern portion of the Delta included branching tributaries and sloughs that lead to the intakes of two large water pumping facilities that divert water from the Delta: the federal Central Valley Project (CVP; D1) and State Water Project (SWP; E1, E2) (Fig.1). The northern extent of the study area followed the mainstem San Joaquin River until its confluence with the Sacramento River just upstream of Chipps Island (A15) and continued through the estuary to the Golden Gate Bridge (A17).

**Figure 1.**
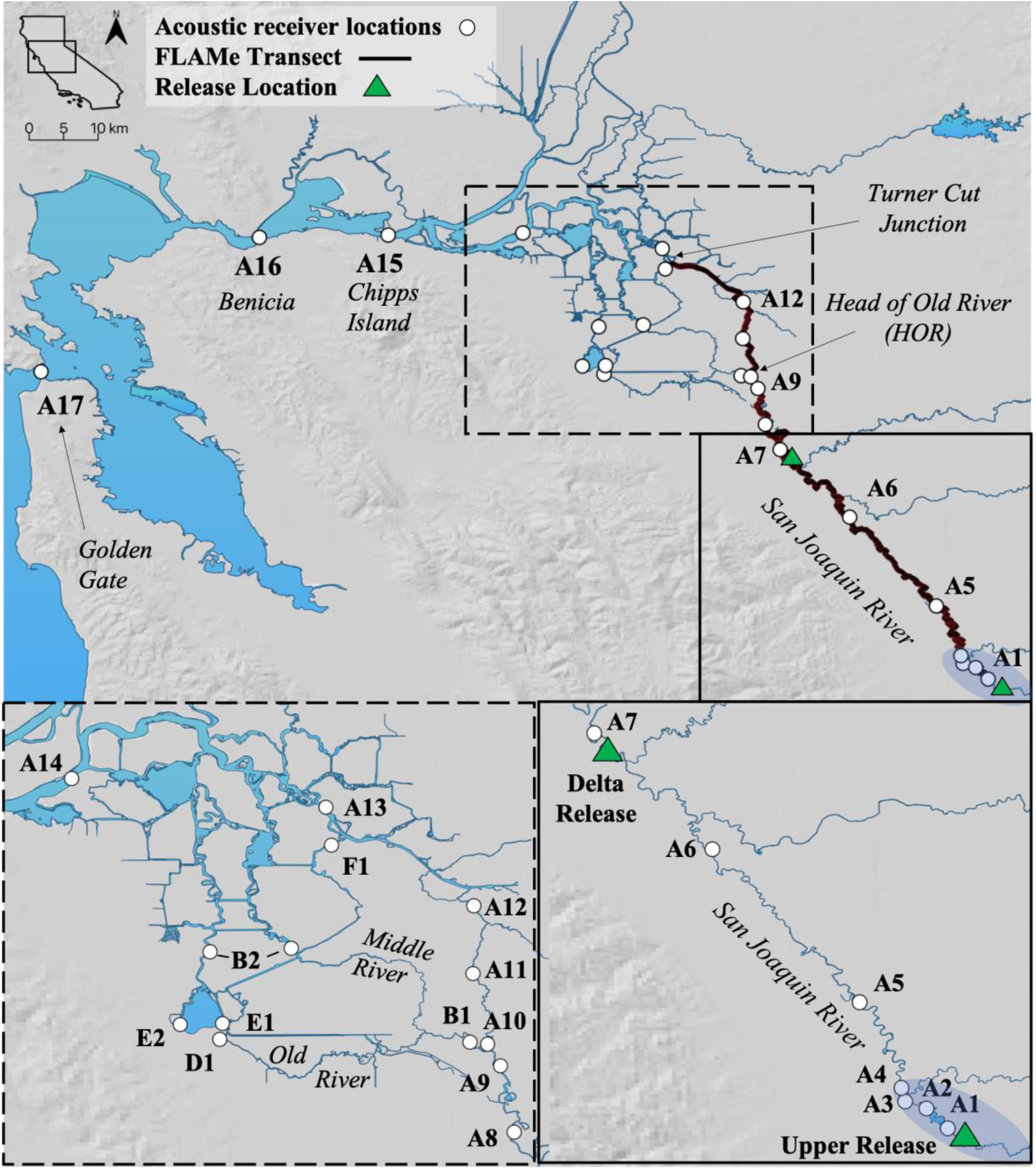
Map of study area from the upper release location at Freemont Ford to the entrance to the Pacific Ocean at the Golden Gate Bridge. White dots mark locations of acoustic telemetry receivers (see Table 1 for additional receiver station information), triangles mark the two fish release locations (upper and Delta releases), the bold black line represents the length of the water quality (FLAMe) transect (n = 3), and the blue shaded area delineates the SJRRP Restoration Area (river only). The two lower panels provide a detailed view of the receiver array within the San Joaquin River (right) and the interior Delta (left). Map created in QGIS 3.6.

**Table 1.** Alphanumeric receiver IDs with their corresponding location name, river kilometer (distance from the Pacific Ocean entrance) and route. *Highway 4 receivers represent the pooled location of Old River and Middle River receivers near the Highway 4 overpass crossings.

| Receiver | Location Name | rkm | Route |
| --- | --- | --- | --- |
| A1 | Below Upper Release | 270 | San Joaquin River |
| A2 | Mud Slough Confluence | 262 | San Joaquin River |
| A3 | Newman | 259 | San Joaquin River |
| A4 | Hills Ferry | 257 | San Joaquin River |
| A5 | Crows Landing | 239 | San Joaquin River |
| A6 | Grayson | 210 | San Joaquin River |
| A7 | Durham Ferry | 181 | San Joaquin River |
| A8 | BCA | 171 | San Joaquin River |
| A9 | Mossdale | 163 | San Joaquin River |
| A10 | SJ HOR (SJ Head of Old River) | 158 | San Joaquin River |
| B1 | OR HOR (OR Head of Old River) | 158 | Old River |
| A11 | Howard | 148 | San Joaquin River |
| D1 | CVP (Central Valley Project) | 144 | Old River |
| E1 | SWP (State Water Project) Forebay | 142 | Old River |
| E2 | SWP (State Water Project) interior channel | 140 | Old River |
| A12 | SJG (SJ Garwood) | 140 | San Joaquin River |
| B2 | Highway 4 Receivers* | 136 | Old River |
| F1 | Turner Cut | 127 | Turner Cut |
| A13 | McDonald Island | 122 | San Joaquin River |
| A14 | Jersey Point | 93 | San Joaquin River |
| A15 | Chipps Island | 71 | All routes |
| A16 | Benicia Bridge | 52 | All routes |
| A17 | Golden Gate Bridge | 1 | All routes |

We investigated two primary migration routes in this study: the mainstem San Joaquin River and the Old River route (Fig. 1, Table 1). Upon reaching the Head of Old River (HOR) junction, fish that remained in the mainstem San Joaquin River were routed through the Port of Stockton (A12) and either migrated through Jersey Point (A14) or entered the interior Delta via Turner Cut (F1) (Fig.1). While this was not the exclusive route for fish into the interior Delta, Chinook salmon have been observed entering Turner Cut in previous studies (Buchanan et al. 2013) and receivers were prioritized at this junction. Fish that deviated from the mainstem San Joaquin River at the HOR entered into Old River, which resulted in a more complex network of routing options (Fig. 1). Here, fish could circumvent the pumping facilities by staying within Old River or Middle River (B2) or enter into either the CVP (D1) or SWP (E1, E2). Fish that survived entrainment in either pumping facility had the opportunity to be salvaged, trucked, and released upstream of Chipps Island (A15) (Fig.1), where all routes converged. Fish that were entrained but not salvaged were treated as mortalities.

### FLAMe

To map habitat heterogeneity along the smolt emigration corridor in the mainstem San Joaquin River, a boat-mounted flow-through sampling system was constructed, modeled after the design of Crawford et al. (2015). While running at high speeds (25-30 km hr^-1^), a ballast pump transported river water through the intake manifold and into an in-line filter to prevent large particles and detritus from compromising sensors. Water was then split at a Y-valve and routed into two separate flow-controlled tanks. Each tank was equipped with one sensor unit, which included a Seabird Scientific Suna V2 optical nitrate (NO_3_) sensor (Bellevue, WA) and a YSI Exo2 multiparameter sonde (Yellow Springs, OH) measuring pH, temperature, DO, turbidity, specific conductivity, chlorophyll-α, and fluorescent dissolved organic matter (fDOM). A Garmin 16x HVS GPS (Olathe, KS) was mounted to the transom of the boat to georeference all measurements. Flow-controlled tanks measured 3 L in volume (including displacement from sensors), and flow rate into each chamber averaged 0.45 L·s^-1^. Sensor measurements were recorded at a frequency of one measurement per second during transects.

Measurements from all three units were recorded in real-time and integrated into one data file via a Campbell Scientific CR 1000 datalogger (Logan, UT). To validate Suna and YSI Exo2 (FLAMe) measurements, we collected discrete water samples at twelve sites along the FLAMe transect for lab analysis of NO_3_, chlorophyll-α, and dissolved organic carbon (DOC, served as a proxy for fDOM). At the same locations, we performed side-by-side measurements (*in situ*) using a YSI 6820 to validate temperature, DO, specific conductivity, turbidity, and pH measurements recorded by the YSI Exo2. Methods for validating measurements collected by the Suna and YSI Exo2 are provided in Supplemental Methods^2^.

FLAMe transects took place at three intervals over the course of juvenile emigration (Table 2, Fig. 2), which, based on previous observations, typically spans from late February/early March through May (Singer et al. in prep^1^). All transects were executed in a downstream direction starting at the upper release site. Transects 2 and 3 ended at the McDonald Island receiver location (A13, Fig.1). Due to time and logistical constraints, transect 1 ended at the HOR junction (Fig.1). Complete FLAMe datasets are provided as open access data files within Dryad Digital Repository (https://doi.org/10.25338/B8VH0D).

**Figure 2.**
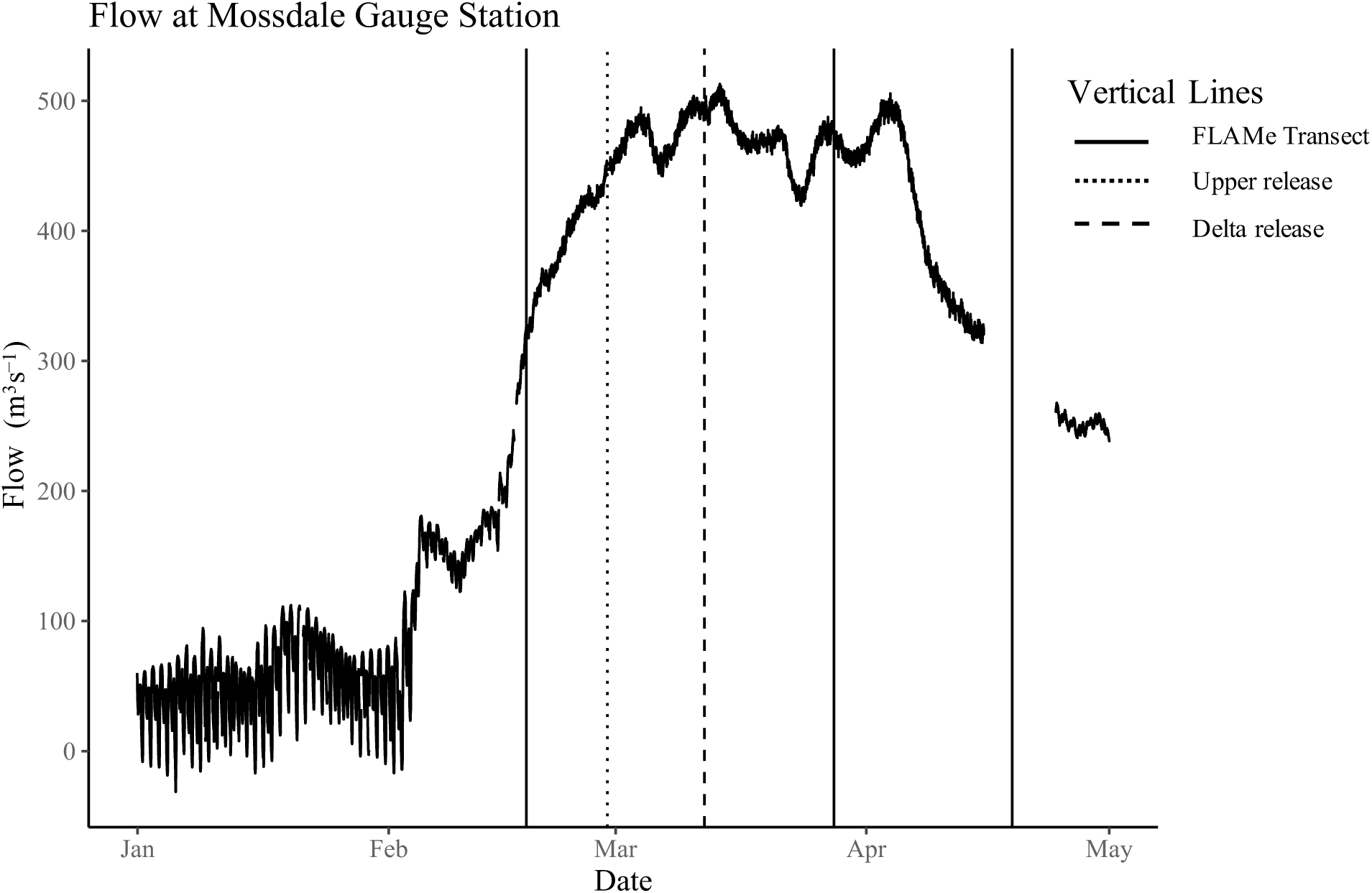
Graph of river flow (m^3^·s^-1^) at the Mossdale CDEC gauge station (station ID = MSD). Solid vertical lines mark the dates when water quality transects were performed, and dotted and dashed vertical lines indicate the timing of upper and Delta fish releases (respectively). Data retrieved from http://cdec.water.ca.gov/.

**Table 2.**
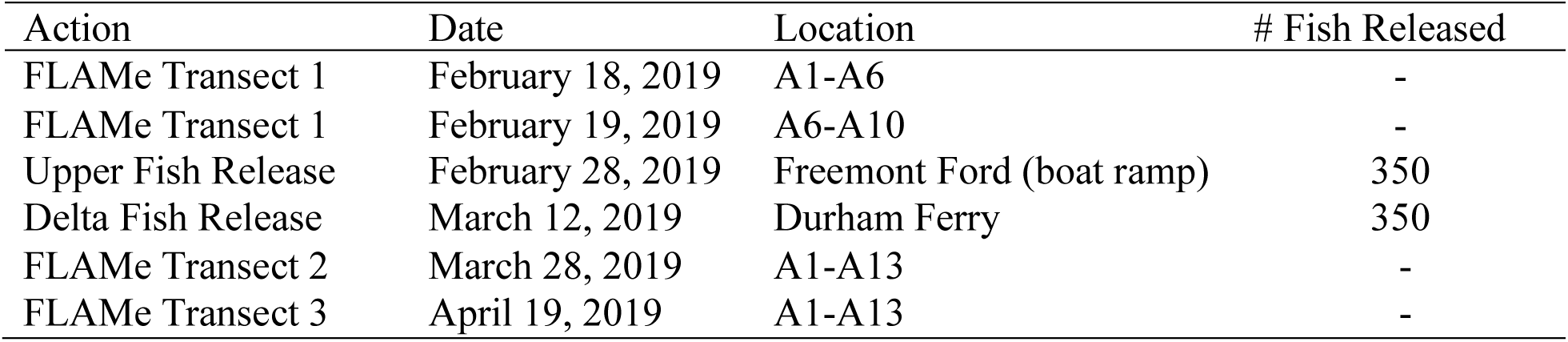
Dates of each FLAMe sampling event and fish release, as well as number of fish released at each site. See Figure 1 for spatial representation of FLAMe sampling and release locations.

### Receiver array, fish tagging, and releases

We supplemented our self-deployed array with acoustic receivers maintained by National Oceanic and Atmospheric Administration (NOAA). In all cases, receivers were deployed as either autonomous (n = 98) or real-time (n = 35) from the lower end of the San Joaquin River Restoration Program (SJRRP) Restoration Area (A1) to the Golden Gate Bridge (A17) (Fig. 1). Receivers spanned routes through the mainstem San Joaquin River, Delta, and estuary, and consisted of Teknologic (Edmonds, WA) and Advanced Telemetry Systems (ATS, Isanti, MN) technologies.

A total of 750 spring-run Chinook Salmon smolts were obtained from the California Department of Fish and Wildlife’s (CDFW) Salmon Conservation and Research Facility (SCARF, Friant, CA; *Section 10(a)(1)(A) permit #1778*) and surgically implanted with JSATS acoustic transmitters following methods similar to those in Singer et al. (2013). In accordance with UC Davis *Institutional Animal Care and Use Committee (IACUC) Protocol Number 21614*, smolts ranging in size from 73-93 mm were anesthetized with a solution of tricaine methanesulfonate (MS-222) (90 mg L^-1^ buffered with 4.32 g sodium bicarbonate) and tags were inserted into the coelom of the fish through a 5-8 mm incision parallel to the mid-ventral line and anterior to the pelvic girdle. Incisions were secured with one suture tied with a 2×2 surgeon’s knot. A total of 350 fish were tagged for each of two release groups and another 50 fish were tagged as part of a concurrent tag effects study. After a minimum 24-hour holding period post-surgery, 350 tagged smolts were released at the upper release site in the SJRRP Restoration Area (A1, Fig.1, Table 2) on February 28, 2019. The second group of 350 fish were released at the Delta release site located at Durham Ferry 12 days after the first release (Fig. 1, Table 2). The purpose of the Delta release was to ensure our ability to estimate survival in the lower reaches of the study area in the event of low survival from the upstream release group. A more detailed description of the surgical and release procedure is provided in Supplemental Methods^2^.

### Statistical Methods

#### FLAMe Data

Raw water quality data from FLAMe surveys were post-processed to remove erroneous readings and blank records, and all data points were snapped to the river centerline using the R package (R Core Team 2017) *riverdist* (v0.15.0; Tyers 2020). Additionally, any GPS coordinate with missing water quality data was removed from the impacted transect (Supplemental Methods^2^). As a result of each transect containing slightly different GPS coordinates and data loss due to post-processing (26%), some coordinates contained values from only one transect, posing a challenge for consistent statistical summaries. Thus, generalized additive mixed models (GAMMs) were used to estimate smooth functional relationships between predictor (distance from upper release location) and response variables (water quality values) (Pedersen et al. 2019). Using the “gam” function and REML smoothing selection method within the R package *mgcv* (v1.8-31; Wood 2011), each water quality variable was modeled across distance to produce one estimate per river kilometer value, consistent across all FLAMe transects. Model results were evaluated using diagnostic tools such as plots of residuals against linear predictors, distribution of residuals, and response against fitted values.

Predicted values from the GAMM analysis were summarized across transects to obtain the mean and coefficient of variation (CV) values for all variables measured at each GPS coordinate and corresponding river kilometer. This ultimately resulted in two primary datasets representing unique characteristics of the environmental data: 1) spatial variation in water quality variables, represented by the mean, and 2) temporal variation in the same variables, represented by the CV of that variable over time. The mean and CV were calculated from n = 3 data points at each coordinate upstream of HOR (where transect 1 ended) and from n = 2 data points at coordinates downstream of HOR. Calculation of the mean across each coordinate integrated over the temporal component of the data, resulting in a dataset representative of variability across space. CV was calculated as the ratio of the standard deviation to the mean (Brown 1998) and resulted in a spatial dataset of variation in environmental variables over time.

#### Classification of habitats along the San Joaquin River

Agglomerative hierarchical cluster (AHC) analysis (a numerical classification method) was used to generate an ecological zonation scheme for the San Joaquin River using the mean (i.e., spatial) and CV (i.e., temporal) environmental datasets, with models following Borcard et al. (2011). Clustering methods were well-suited for our application as they do not require *a priori* knowledge of classification groups (Yohannes and Webb 1999; Safavian and Landgrebe 1991). While many types of clustering methods exist, hierarchical methods are well-developed, dynamic, and widely used in ecological research (Boesch 1977).

A Euclidean distance matrix was calculated for each log-transformed and standardized (z-score normalization) dataset, which was then partitioned into hierarchical clusters via the Ward method. The number of interpretable clusters was determined using the silhouette method as implemented in the R package *cluster* (v2.1.0; Maechler et al. 2019), and cluster stability was confirmed using bootstrap sampling in the “clusterboot” function of the R package *fpc* (v2.2-5; Hennig 2020). Once the number of clusters was determined for each dataset, the spatial extent of each cluster was adjusted to fit within the bounds of the nearest receiver detection sites.

Following AHC analysis, we performed Principle Component Analysis (PCA) to identify the most meaningful variables in the dataset while minimizing loss of original information (Helena et al. 2000). In reducing dimensionality to the axes (i.e. “principle components”) that hold the most information, we identified key variables that characterized the most variation in each dataset (Gotelli and Ellison 2004). Axes with eigenvalues (measures of axis variance) greater than the mean of all eigenvalues were retained for analysis, as defined by the Kaiser Guttman criterion (Borcard et al. 2011). We applied the broken-stick model following Peres-Neto et al. (2003) to assess the degree of association of variables with ordination axes (Frontier 1976; Borcard et al. 2011).

#### Acoustic Telemetry Data

Raw acoustic telemetry data files were processed to identify and remove false positive and multipath detections using a filtering algorithm adapted from the University of Washington Columbia Basin Research Group. Detections were analyzed for validity based on three or more occurrences of a tag at the estimated nominal pulse rate interval (PRI, 5 s for this study) within a 12-interval (i.e., 60-s) rolling window. A more detailed definition of filter criteria can be found at www.cbr.washington.edu/analysis/apps/fast. Additionally, we applied a behavior-based predator filter that evaluated movement patterns for each tag according to assumptions of differences between predator and smolt movement patterns (Vogel 2010; Buchanan et al. 2013; Singer et al. in prep^1^). Additional details on the acoustic telemetry data processing procedure and detection history formation are available in Supplemental Methods^2^.

A multistate release-recapture model was fit to the data using maximum likelihood estimation in the software program USER (Lady and Skalski 2009) following methods similar to those in Buchanan et al. (2013) and Singer et al. (2020 in prep^1^). In brief, each “state” in the multistate model structure represents a pathway through the Delta, and the probability of observing each detection history becomes a function of the following parameters: survival (*S*), probability of detection at a receiver location (*p*), route selection (*ψ*), and transition probability (*ϕ*) (Fig. S1). Transition probability (*ϕ*) is defined as the joint probability of movement towards a particular route and survival along that route (*ψS*) and was used in cases where survival and route entrainment probabilities could not be separately estimated (Perry et al. 2010; Buchanan et al. 2013). Due to sparse detections at Middle River (n = 2 individuals), the Middle River and Old River receivers (B2, Fig.1) were pooled (hereafter, “Highway 4 Receivers”; Table 1) and *ϕ*_*B*1*B*2_became the total transition probability from the HOR (B1) to either Highway 4 location. Compared to the upper release group, there was the possibility that newly released Delta fish had different probabilities of survival, migration, and detection through the initial reaches downstream of the Durham Ferry release site (Fig. 1). Therefore, separate models were initially constructed with unique parameters for each release group and then tested against simpler models with common parameters between the two release groups (Buchanan et al. 2013). This procedure was followed until a final model was built maximizing the number of common parameters without reducing model fit, as determined by Akaike’s Information Criterion (AIC) (Burnham and Anderson 2004).

#### Candidate Survival Models

We developed four models (Table 3) representing competing hypotheses on the relationship between outmigration survival and regional habitat characteristics:

1. Survival is a function of distance travelled, independent of habitat characteristics (H_0_)
2. Survival is a function of unknown reach-specific variables (H_1_)
3. Survival is a function of environmental gradients, informed by water chemistry variables that either vary over (a) space (H_2_) or (b) time (H_3_)

**Table 3.** Multistate mark-recapture model results for competing hypotheses on spatial variation in survival.

| Model | Description | Hypothesis | $\Delta$ AIC |
| --- | --- | --- | --- |
| M<sub>0</sub> | Common-survival rate model | Survival is a function of distance alone, and thus information provided by regional differences is minimal | 637 |
| M<sub>1</sub> | Geographic model | Survival is a function of unknown site-specific variables contained within each reach of the array | 13 |
| M<sub>2</sub> | Spatial model | Survival is a function of ecologically distinct regions that vary *spatially* | 0 |
| M<sub>3</sub> | Temporal model | Survival is a function of ecologically distinct regions that vary *temporally* | 132 |

The goal was to determine if variation in survival rate (survival per km) through the 150 km mainstem section of river sampled by the FLAMe (A1-A15, Fig.1) could be explained by regional differences in habitat. A “geographic model” (M_1_) represented the most complex parameterization of survival, with survival rate (per km) estimated uniquely for each reach (i.e., the area between two receivers) within the receiver array (H_1_). The other three models were compared to the geographic model.

The first alternative model considered represented the hypothesis that survival was a function of distance alone (H_0_), and thus any information provided by regional differences was minimal (M_0_). Therefore, this model assumed a common survival rate for all the reaches and regions in the study area. The final two models reflected ecologically distinct regions in the river resulting from AHC analysis on the water quality datasets. The spatial model (M_2_) grouped reaches based on the spatial distribution of the clusters identified from the mean of the water quality transects, and reflected the hypothesis that survival is a function of ecologically distinct zones that vary across space (H_2_). The temporal model (M_3_) grouped reaches based on the spatial distribution of the clusters identified from the CV of the transects, which represented the hypothesis that survival is a function of ecologically distinct zones that vary over time (H_3_). Model selection based on AIC was used to identify the hypothesis best supported by the data.

#### Alternative parameterizations of reach-specific survival (*S*_*i*_)

The same underlying model structure of *S*, *p*, *ψ* and *ϕ* parameters was used in all four models (Table 3), with *S* parameterized in terms of the per-km survival rate *σ* such that *S* = *σ*^*d*^, where *d* = reach distance (km). However, parameterization of *σ* and *ϕ* differed for each model depending on the hypothesis tested. For the geographic model (M_1_), probability of survival *S* through reach *i* was defined as 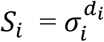 where *S* = probability of survival, *σ* = per-km survival rate, *d* = distance (km), and *i* = reach. This resulted in a unique estimate of *σ* for each reach within the array. Additionally, regional survival probabilities were estimated, defined as functions of route entrainment probabilities (*ψ*) and route-specific survival probabilities (*S*) (Perry et al. 2010; Buchanan et al. 2013). For example, survival through the Delta via the mainstem San Joaquin River route was defined as:

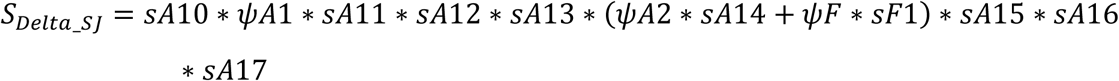

where *sAi* is the probability of survival from site *i*-1 to site *i*, conditional on survival to site *i*-1 (Fig. S1).

For the distance-based model (M_0_), probability of survival *S*through reach *i*was defined as *S*_*i*_= *σ*^*di*^, resulting in a single estimate of *σ* throughout the array. This required modeling both the reach-specific survival *S*_*i*_and transition probabilities *ϕ*_*ij*_in terms of a single survival rate, which allowed for survival to be estimated independently of route selection at the junctions where this was not possible in the competing models (M_1_, M_2_, M_3_). Therefore, we redefined the transition probability *ϕ* for this model as *ϕ*_*ij*_= *σ*^*di*^*ψ*_*j*_, where *ϕ*_*ij*_= transition probability from site *i* to site *j*, *σ* = common per-km survival rate, *d*_*i*_= distance (km) from site *i* to site *j*, and *ψ*_*j*_ = probability of entrainment in route *j*.

For the cluster models (M_2_ and M_3_), separate *σ*parameters were estimated for each cluster, as defined by the AHC analysis. For example, if reaches 1-4 were contained within cluster 1, the total probability of survival through this region was defined as *S*_*C*1_ = 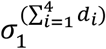 where *S*_*C*1_ = survival probability through cluster 1, *σ*_1_ = survival rate per km in cluster 1, and 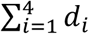 is the sum of the lengths (km) of reaches 1-4. In mapping the clusters, we identified the receiver locations nearest cluster boundaries to enable estimation of the survival rate parameter *σ* in each zone. Survival in reaches excluded from the FLAMe transects was parameterized as in model M_1_.

## Results

### Water Quality Transects

According to CDEC Water Year Hydrologic Classification Indices, 2019 was considered a ‘wet’ year in the San Joaquin Basin (http://cdec.water.ca.gov). Water quality sampling took place at three intervals over the course of the juvenile emigration window (Fig. 2), during which time flows were higher than average due to heavy rains at the start of the sampling period. Regression analysis of lab-analyzed/YSI 6820 and FLAMe measurements generally reflected strong linear relationships, suggesting accurate characterization of water chemistry variables by the FLAMe (Fig. S2, Supplemental Results^2^).

Total number of observations for all transects used in the GAMM analysis was 20,183, and represented 74% of the original dataset after data cleaning (Fig. S3a-c). Raw FLAMe measurements were in close agreement to GAMM-predicted values of environmental data (Fig. S4). From the AHC analysis, silhouette width suggested seven unique clusters for the spatial dataset (i.e., mean values) and three for the temporal dataset (i.e., CV values). Bootstrap sampling of the original data matrices confirmed three clusters for the temporal dataset, but resulted in higher stability indices for six clusters in the spatial dataset. Therefore, six clusters were ultimately used in analysis of the spatial dataset. Because the receiver array was predetermined, some cluster boundaries were jittered to fit within reaches (adjustment ranging from 0.15-9.1 km) (Fig. 3). Cluster results from the spatial dataset indicated six distinct zones along the longitudinal profile of the river: the restoration area (C1), upper river (C2), mid-river (C3), lower river (C4), southern Delta (C5), and mainstem Central Delta (C6) (Fig. 3A). The spatial distribution of cluster results from the temporal dataset indicated three regions with distinct differences in temporal variation: upper river (C1), lower river (C2), and Delta (C3) (Fig. 3B).

**Figure 3.**
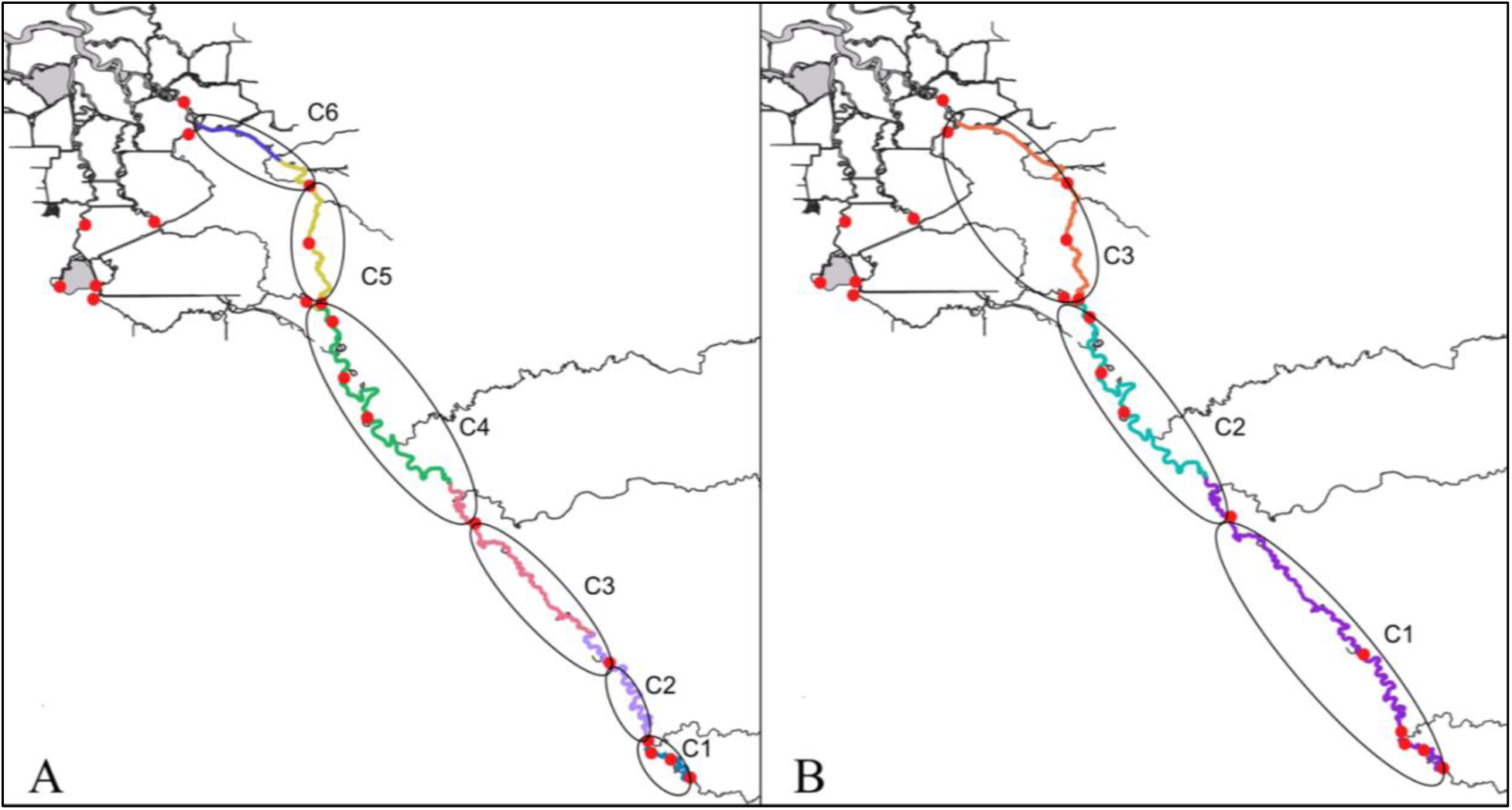
Spatial distribution of cluster groups (C1-C6) derived from AHC analysis on the mean (panel A) and CV (panel B) values of water quality parameters across the three transects.

Environmental conditions in the spatial dataset (i.e., mean values) exhibited directional trends across clusters starting from the restoration area (C1, Fig. 3A) and moving downstream to the Central Delta (C6, Fig. 3A), with the exception of temperature, NO_3_, and pH (Table S1, Fig. 4). The highest values of chlorophyll-α, specific conductivity, temperature, fDOM, and turbidity characterized the restoration area (C1, Fig. 3A) (Table S1, Fig.4). Values of these five variables decreased from the restoration area (C1) to the lower river (C4), after which only temperature increased through the southern Delta (C5). Minimum values of chlorophyll-*α* and turbidity occurred in the mainstem Central Delta (C6), while temperatures continued to increase in this region. Dissolved oxygen followed an opposite trend, where values were lowest in the restoration area (C1) and gradually increased downriver (C2-C4), ultimately peaking in the Delta regions (C5-C6) (Table S1, Fig. 4). The Kaiser Guttman criterion confirmed two principal components (PCs) that explained 86% of the total variance in the spatial dataset. Accounting for 67% of the total variance, the first PC was not significantly correlated with any of the environmental variables according to the broken-stick model (Table S2a). The second PC, accounting for 19% of the total variance, was correlated with temperature (Table S2a).

**Figure 4.**
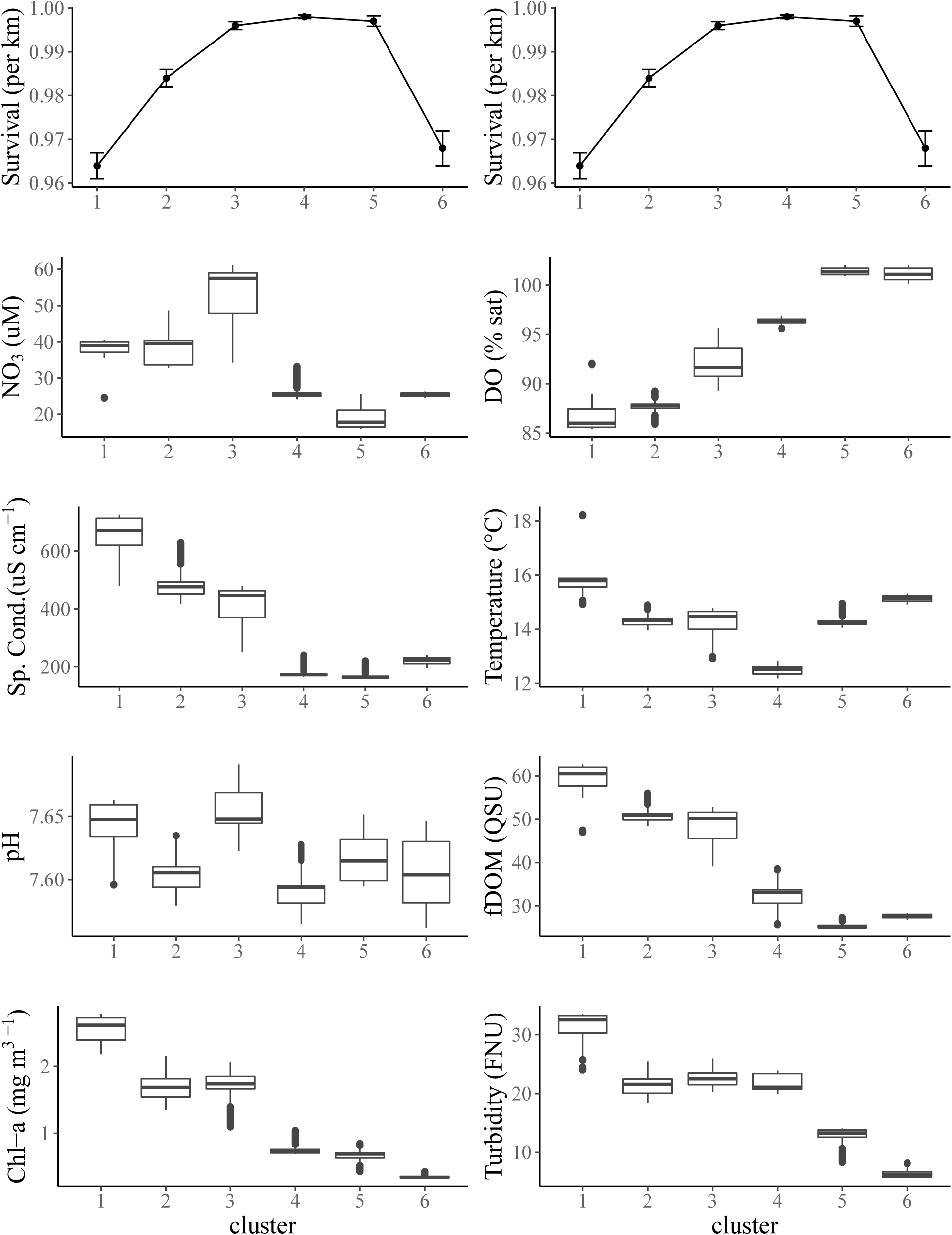
Water chemistry distributions by variable and cluster in relation to per km survival rate. Cluster groupings were derived from AHC analysis on the spatial dataset, and survival rates were estimated from model M_2_. The bold horizontal lines represent median values, while the upper and lower edges of the boxes represent the 75^th^ (Q3) and 25^th^ (Q1) percentiles, respectively. The upper and lower ends of the vertical lines represent largest and smallest value no further than 1.5 *IQR (interquartile range, or Q3-Q1) from the 75^th^ and 25^th^ quartiles. Points beyond the end of the vertical lines represent outliers. Note: The top two graphs of survival rate are identical plots.

Temporal variation of water quality data across the three clusters (i.e., dataset of CV values) was generally lowest in the Delta for all environmental variables (C3, Fig. 3B), with the exception of chlorophyll-*α* and pH (Fig. S5). Chlorophyll-*α* and specific conductivity had the greatest variation in the upper river (C1) (median CV = 0.67 and 0.52, respectively), while NO_3_ and turbidity were most variable across time in the lower river (C2) (median CV = 0.81 and 0.71, respectively) (Fig. 3B, Fig. S5). The Kaiser Guttman criterion confirmed two PCs that explained 86% of the total variance in the temporal dataset, 47% accounted for by the first PC and 39% by the second PC (Table S2b). According to the broken-stick model, temperature and fDOM were correlated with the first and second PCs, respectively (Table S2b).

### Model Selection and Survival Estimates

The spatial model (M_2_) was selected by AIC over all alternative models (Table 3) as the best representation of spatial differences in smolt survival. However, the comparatively low ΔAIC value of 13 also illustrated support for the geographic model (M_1_), which out-performed both the temporal (M_3_) and the common survival-rate model (M_0_) (Table 3). Survival estimates are hereafter presented as two metrics: (1) survival per-km 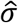 (for comparison between reaches/clusters) and (2) total survival probability through a reach/cluster, represented by 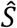. Both terms refer to the same reach or cluster (cluster = a group of reaches, see Table S3). In the spatial model (M_2_), the estimated per-km survival rate was low through the restoration area (C1) (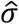 = 0.964, 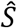 = 0.61 ± 0.02 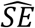) (Fig.3A, Fig.4). Survival rates increased through the next four sections of river, estimated at 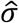 = 0.984 (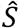 = 0.76 ± 0.03 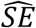), 0.996 (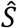 = 0.89 ± 0.02 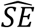), 0.998 (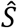 = 0.94 ± 0.02 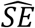), and 0.998 (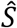 = 0.96 ± 0.02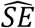) (clusters C2-C5 respectively, Table S3). The probability that fish remained in the mainstem San Joaquin River upon reaching the HOR junction was estimated at 0.59 (± 0.04 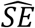). Survival rate decreased considerably through the mainstem Central Delta (C6), the final region through which environmental measurements were recorded (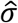 = 0.968, 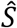 = 0.63 ± 0.04 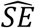) (Fig. 3A, Fig. 4) (Table S3).

Survival estimates through reaches excluded in the FLAMe transect were calculated from the geographic model (M_1,_ Table 3). Survival (*S*) from the Turner Cut junction to Chipps Island (A15) (through both the mainstem river (A14) and Turner Cut (F1) pathways) was 0.68 (± 0.02 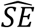) (Table S3). Of the fish that selected Old River at the HOR junction, more than half were estimated to enter into the SWP (E1) forebay (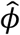 = 0.58 ± 0.07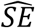). Of these fish, very few survived to Chipps Island (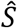 = 0.18 ± 0.06 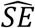). Overall, survival from Old River to Chipps Island was estimated at 0.076 (± 0.03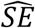). Regional survival from Chipps Island (where all routes converged) to the Golden Gate Bridge (A17) was 0.95 (± 0.03 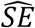). The estimated cumulative probability of survival through the entire study area, from the upper release location to the Golden Gate Bridge (A17), was 0.05 (± 0.009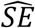) (Table S3). No adjustments were made to survival estimates due to premature transmitter failure, excessive tag shedding, or tag-induced mortality (See Supplemental Results^2^ for additional details on tag effects).

## Discussion

Successful reintroduction and management of salmonids in the Central Valley necessitates an improved understanding of the habitat conditions driving population level success at every life stage (Zeug et al. 2019). Previous work investigating smolt outmigration in the San Joaquin River has focused on spatiotemporal patterns in mortality (Buchanan et al. 2013, 2018; Singer et al. in prep^1^), which is an important step towards informing actionable management (Perry et al. 2010). However, many of these studies highlight a limited understanding of how spatially-explicit habitat conditions shape survival patterns. In the present study, we used a multiscale approach towards understanding relationships between outmigration survival and localized habitat variation by employing a novel application of limnological technology (FLAMe). Our analysis of environmental gradients across a 150 km section of the mainstem San Joaquin River revealed that variation in survival was better explained by water chemistry conditions that varied over space (M_2_) than conditions that varied over time (M_3_) (Table 3). Furthermore, this study found that continuous, high-resolution spatial data can be used to identify habitat gradients that hold more information about survival dynamics than the commonly used geographic model (M_1_), thus providing ecological context to salmon survival throughout the river.

Survival of juvenile spring-run Chinook Salmon was estimated to be lowest in two regions within the longitudinal transect, the restoration area (C1) and the mainstem Central Delta (C6) (Fig. 3A, Fig. 4). The physical landscape of the restoration area (C1) is more representative of a natural environment than the remaining downstream reaches. The restoration area is dominated by salt basin and high marsh vegetation, and minimal levee infrastructure enables lateral connectivity to the riparian edge during flood conditions (such as those that occurred in 2019) (SJRRP 2019). This floodplain connectivity mediates delivery of sediments, nutrients, and organic materials to the river (Bisson et al. 1987), evident as higher levels of allochthonous dissolved organic matter (i.e., higher fDOM concentrations) (Fig. 4). Organic matter inputs are vital energy sources for most lotic and riverine food webs (Hall et al. 2000), and likely contributed to primary productivity in this region. Chlorophyll-*α* values measured in the restoration area (C1) were the highest recorded across all three transects (Fig. S4). Carbon-rich habitats enhance secondary production, which can provide additional crucial food resources for migrating juvenile salmon (Claeson et al. 2006; Jeffres et al. 2020). Additionally, high turbidity levels measured in this region could aid juvenile salmon in predator avoidance, as well as decrease predator encounter rates by increasing migration rate (Gregory and Levings 1998; Michel et al. 2015).

Despite exhibiting characteristics of high-quality juvenile salmon habitat, the restoration area (C1) reflected the lowest survival rates estimated throughout the FLAMe transect (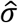 = 0.964, 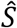 = 0.61 ± 0.02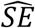). Mean temperature values were highest in this region compared to the rest of the transect during all three sampling events, suggesting that this area remained relatively warmer throughout the migration period (February-April). All tagged salmon were downstream of this region when temperatures reached the upper thermal ranges (> 18 °C, outliers in Fig. 4), and therefore the interquartile range (15.5 °C - 15.8 °C, Fig. 4) is likely more reflective of what the majority of fish were experiencing in the restoration area (C1). While these temperatures may not be lethal, increased temperatures can negatively affect the overall ecology of many fishes, but especially coldwater species (Magnuson et al. 1979; Lyons et al. 2009). For example, temperature can influence body size and spatial distribution of warmwater predators (Rypel 2014). Catch data from fyke nets and electrofishing surveys in the restoration area (C1) have recorded high numbers of non-native piscivores, including known salmonid predators such as such as channel catfish (*Ictalurus punctanus*), largemouth bass (*Micropterus salmonides*), and striped bass (*Morone Saxatilis*) (Portz et al. 2013; Grossman et al. 2013). Previous research found predation risk on juvenile salmon in the Delta is positively correlated with temperature (Michel et al. 2019), which could pose problems for smolts upstream if similar temperature-predation relationships exist in riverine environments.

The second major mortality sink along the water quality transect occurred in the mainstem Central Delta (C6) (Fig. 3A), which was characterized by low turbidity, chlorophyll-*α*, and fDOM values, as well as warmer temperatures (Fig. 4). High water clarity can be problematic for juvenile salmon, as turbidity has proven to be a key environmental factor for predation evasion by decreasing the visual field of predators (Gregory and Levings 1998). Dams, diversions, and levees contribute to declines in turbidity by choking off sources of suspended sediments to the Delta (Weitkamp 1994). This effect is exacerbated by overgrowth of non-native aquatic macrophytes, as vegetation promotes sediment deposition (thus decreasing turbidity) by increasing vertical drag and decreasing flow in the water column (Grossman et al. 2013). The reduction in incidence and risk of piscivory provided by turbid conditions is important for behavioral processes such as migration (Ginetz and Larkin 1976) and feeding activity (Gregory 1993). Turbid conditions become an important defense against open-water predators like largemouth bass (*Micropterus salmoides*) (Ferrari et al. 2014), as two-dimensional tracking of juvenile Chinook Salmon in the Sacramento River has suggested that outmigrating smolts primarily travel within the center channel (Sandstrom et al. 2013).

In conjunction with high predation risk in the mainstem Central Delta region (C6), food limitation may also influence smolt survival rates. Chlorophyll-*α* measurements in this region were < 1 ug L^-1^ (Fig. 4), consistent with long-term records of diminishing productivity in the Delta (Robinson et al. 2016). Current estimates of phytoplankton production, which form the dominant food supply to primary consumers (such as zooplankton), puts Delta productivity in the lowest 15% of the world’s estuaries (Cloern et al. 2014; Robinson et al. 2016). Parallel declines in primary production and zooplankton abundance in the Delta suggest processes limiting primary production in turn regulate the capacity of these habitats to sustain pelagic food webs (Lucas et al. 2002). Dampened productivity is one of the leading factors hypothesized to contribute to the Pelagic Organism Decline (POD), coined from the observation of major declines in four historically abundant pelagic species in the upper San Francisco estuary over the last two decades (Sommer et al. 2007). Organic matter transferred from upland ecosystems serves as the basis of the aquatic food web (Helfield and Naiman 2001) and loss of lateral connectivity to historically inundated floodplains could be a key factor in observed chlorophyll-*α* and fDOM differences between the restoration area (C1) and mainstem Central Delta habitats (C6).

Higher temperatures were a common factor in the two zones with lowest survival (C1, C6; Fig.3A) and a statistically significant source of variation based on PCA analysis (Table S2). These results underscore the importance of managing habitat for temperature, which is supported by extensive literature on the impacts of temperature on salmon physiology, behavior, and predator bioenergetics, and is likely to be of increased concern under climate change (McCullough et al. 2001; Petersen and Kitchell 2001; Zillig et al. 2021). However, temperature is unlikely to be the sole factor driving similar survival estimates in the restoration area (C1) and mainstem Central Delta (C6). Differences in survival rates between these habitats (C1, C6, Fig. 3A) and the intervening regions (C2-C5, Fig. 3A) were large (estimated at 0.034 difference in per km survival rate, or a 0.33 difference in survival probability) relative to the median difference in temperature (3.27 °C). Relatively small temperature differences between regions with large discrepancies in survival suggest additional factors may be contributing to observed mortality. Low survival in the mainstem central Delta (C6) was likely a result of warm temperatures combined with low food availability (as indicated by low chlorophyll-*α* and fDOM levels) and high exposure to predators (due to high water clarity and little access to predator refugia). In contrast, as previously described, the restoration area (C1) was generally reflective of more complex, higher-quality habitat. Despite favorable conditions, a combination of low DO and warmer temperatures may have interacted to create challenging conditions for newly released hatchery fish. Relatively mild environmental stressors may have been amplified when combined with trucking and handling stress experienced by smolts released into a new environment (Schreck et al. 1989). Therefore, future studies relating salmon survival to habitat characteristics might attempt to account for acclimation dynamics.

Current methods of exploring variation in survival have either characterized survival exclusively from a geographic standpoint (Perry et al. 2010; Buchanan et al. 2013, 2018; Singer et al. in prep^1^) or incorporated environmental covariates that are temporally explicit (Perry et al. 2018; Buchanan and Skalski 2020). These approaches are useful for understanding impacts of covariates that are highly variable through time (e.g. flow) or dominate regional characteristics (e.g. export rates) (Buchanan and Skalski 2020), but may be limited in their ability to describe effects of localized environmental conditions on survival. Our study complements current methods by allowing multiscale habitat data to elucidate environmental gradients that have been previously hypothesized from spatial patterns in survival. Providing ecological context to regional differences in survival is critical for crafting effective management strategies. For example, similar survival rates in the restoration area (C1) and mainstem Central Delta (C6) superficially suggest that common factors may drive survival in these contrasting habitats. Failing to consider spatially-explicit information on the environmental conditions within these regions may lead to faulty assumptions about the factors shaping outmigration survival. Using the information gained from this study, we recommend efforts that improve habitat complexity within the central Delta, specifically those that increase turbidity levels. Prioritizing flows to meet the thermal needs of juvenile salmon throughout their outmigration may have the added benefit of helping address turbidity issues in the Delta as well as improve dissolved oxygen levels in the restoration area.

Ecological classification frameworks are important tools for understanding the scale-dependent processes that govern landscape patterns and organism responses (Wiens 2002; Higgins et al. 2005; Rypel et al. 2019). The FLAMe system provided an empirical method for delineating longitudinal heterogeneity at a scale that could be related to outmigration survival, thus providing ecological context to spatial patterns in mortality. Beyond a telemetry application, this method may serve as a simple, fast way to identify geographic zones within which fisheries management policies or restoration projects could be implemented. Applications of classification tools are many and include improved setting of regional fisheries expectations, use in developing more effective fish stocking rates or harvest regulations, and improved design of scientific studies and monitoring efforts (Rypel et al. 2019; Schupp 1992; Wehrly et al. 2012). Managers could use a classification scheme based on regional patterns to set water quality and fisheries standards that balance human impacts and biological requirements, inform monitoring sites based on major change points within a river, and predict the impacts of land use and pollution controls.

Restoring degraded freshwater ecosystems in the face of global environmental change is a central challenge in fisheries science and ecology (Pahl-Wostl 2007; Davies 2010). Rising to meet this challenge will require novel tools to identify functional relationships between ecosystem processes and species response (Jackson et al. 2016; Van den Brink et al. 2016). Our method of habitat classification follows the general concept that rivers are spatially organized by hierarchically-related physical and ecological processes (Frissell et al. 1986; Thorp et al. 2008), and can therefore be applied to a range of ecological questions throughout large rivers globally. This approach could be useful in understanding the role of alternative stable states in maintaining river habitat complexity (Wohl et al. 2015; Adams 1997), or to identify spatial patterns in early-warning signals of ecological tipping points following habitat alteration (Butitta et al. 2017). While there are many potential applications related to declining fisheries such as native salmonids, these techniques could also be applied to almost any mobile fish species that experiences changes in habitat conditions over sensitive portions of its life cycle. Integrating simple environmental classification frameworks that improve understanding of physical and biological processes will be critical for informed decision-making in natural resource and fisheries management.

## Supporting information

Supplementary Materials for: Survival of a threatened salmon is linked to spatial variability in river conditions

## Acknowledgements

Funding for this study was provided by the Delta Stewardship Council (Award 1469) and the California Agricultural Experimental Station of the University of California Davis, grant number CA-D-WFB-2467-H to ALR and CA-D-WFB-2098-H to NAF. ALR was supported by the Peter B. Moyle & California Trout Endowment for Coldwater Fish Conservation. A special thanks to Rypel and Fangue lab members Wilson Xiong, Amanda Agosta, Mackenzie Miner, Heather Bell, Michael Thomas, and Matthew Pagel for your tireless work in the field and lab, and to Dr. Luke Loken for assistance with FLAMe design and data processing. This work would not have been possible without the help from staff at CDFW Salmon Conservation and Research Facility, SJRRP Fisheries Management Work Group, DWR SWP Fish Facilities Unit, US Bureau of Reclamation, and NOAA for coordinating tagging operations, providing the research subjects, assisting with deployments, granting access to field sites, and sharing telemetry data.

## Contributors Statement

CLH performed project administration, conducted the investigation, managed data curation, performed the formal analysis and data visualization, and led the writing of the original draft. GPS provided supervision, contributed to conceptualization, methodology, and software, and assisted with writing (review and editing). RAB contributed to conceptualization, methodology and analysis, and assisted with writing (review and editing). DEC performed project administration and provided supervision and resources. NAF co-led the funding acquisition, performed project administration, provided resources, and assisted with writing (review and editing). ALR co-led the funding acquisition, performed project administration, provided resources, contributed to conceptualization and methodology, and assisted with writing (review and editing).

## Data Availability

Data are available in Dryad Digital Repository (https://doi.org/10.25338/B8VH0D).

Additionally, data are available in a shiny application web framework for visualization and download (link: http://cftc.metro.ucdavis.edu/biotelemetry-autonomous-real-time-database/resource).

## Footnotes

1 Singer, G.P., C.L. Hause, E.D. Chapman, M.P. Pagel, A.P. Klimley, N.A. Fangue, and A.L. Rypel. Dynamics of juvenile survival for reintroduced spring-run Chinook Salmon smolts. *Manuscript in Preparation*.

2 [See Supplemental Materials]

## References

Adams, W.M. 1997. Rationalization and Conservation: Ecology and the Management of Nature in the United Kingdom. Trans. Inst. Br. Geogr. 22(3):277–291.

Bisson, P.A., R.E. Bilby, M.D. Bryant, C.A. Dolloff, G.B. Grette, R.A. House, M.L. Murphy, K.V. Koski, and J.R. Sedell. 1987. Large woody debris in forested streams in the Pacific Northwest: past, present, and future. *In* Streamside management: forestry and fishery interactions. Edited by E.O. Salo and T.W. Cundy. College of Forest Resources, University of Washington, Seattle, Washington. Contrib. No. 57, pp. 143–190.

Boesch, D.F. 1977. Application of numerical classification in ecological investigations of water pollution. Virginia Institute of Marine Science, Special Scientific Report No. 77. EPA-600/3-77-033.

Borcard, D., F. Gillet, and P. Legendre. 2011. Numerical Ecology with R, Springer Science + Business Media LLC.

Brown, C.E. 1998. Coefficient of Variation. *In* Applied Multivariate Statistics in Geohydrology and Related Sciences. Springer, Berlin, Heidelberg. pp. 155–157.

Buchanan, R.A., J.R. Skalski, P.L. Brandes, and A. Fuller. 2013. Route Use and Survival of Juvenile Chinook Salmon through the San Joaquin River Delta. N. Am. J. Fish. Manage. 33(1):216–229.

Buchanan, R.A., P.L. Brandes, and J.R. Skalski. 2018. Survival of Juvenile Fall-Run Chinook Salmon through the San Joaquin River Delta, California, 2010–2015. N. Am. J. Fish. Manage. 38(3):663–679.

Buchanan, R.A. and J.R. Skalski. 2020. Relating survival of fall-run Chinook Salmon through the San Joaquin Delta to river flow. Environ. Biol. Fishes. 103:389–410.

Burnham, K.P. and D.R. Anderson. 2004. Multimodel Inference: Understanding AIC and BIC in Model Selection. Sociol. Methods Res. 33(2):261–304.

Butitta, V.L., S.R. Carpenter, L.C. Loken, M.L. Pace, and E.H. Stanley. 2017. Spatial early warning signals in a lake manipulation. Ecosphere, 8(10).

Claeson, S.M., J.L. Li, J.E. Compton, and P.A. Bisson. 2006. Response of nutrients, biofilm, and benthic insects to salmon carcass addition. Can. J. Fish. Aquat. Sci. 63(6):1230–1241.

Clark, W.C, J.E. Shelbourn, and J.R. Brett. 1981. Effect of artificial photoperiod cycles, temperature, and salinity on growth and smolting in underling coho (*Onchorhyncus kisutch*), chinook (*O. Tshawytscha*), and sockeye (*O. nerka*) salmon. Aquaculture, 22:105–116.

Cloern, J.E., S.Q. Foster, and A.E. Kleckner. 2014. Phytoplankton primary production in the world’s estuarine-coastal ecosystems. Biogeosciences, 11:2477–2501.

Crawford, J.T., L.C. Loken, N.J. Casson, C. Smith, A.G. Stone, and L.A. Winslow. 2015. High-Speed Limnology: Using Advanced Sensors to Investigate Spatial Variability in Biogeochemistry and Hydrology. Environ. Sci. Technol. 49(1):442–450.

Davies, P.M. 2010. Climate Change Implications for River Restoration in Global Biodiversity Hotspots. Restor. Ecol. 18(3):261–268.

Fausch, K.D., C.E. Torgersen, C.V. Baxter, and H.W. Li. 2002. Landscapes to Riverscapes: Bridging the Gap between Research and Conservation of Stream Fishes: A Continuous View of the River is Needed to Understand How Processes Interacting among Scales Set the Context for Stream Fishes and Their Habitat. BioScience, 52(6):483–498.

Ferrari, M.C.O., L. Ranåker, K.L Weinersmith, M.J. Young, A. Sih, and J.L Conrad. 2014. Effects of turbidity and an invasive waterweed on predation by introduced largemouth bass. Environ. Biol. Fishes, 97(1): 79–90. doi:10.1007/s10641-013-0125-7.

Fisher, F.W. 1994. Past and Present Status of Central Valley Chinook Salmon. Conserv. Biol. 8(3):870–873.

Frissell, C.A., W.J. Liss, C.E. Warren, and M.D. Hurley. 1986. A hierarchical framework for stream habitat classification: Viewing streams in a watershed context. Environ. Manage. 10(2):199–214.

Frontier, S. 1976. Etude de la decroissance des valeurs propes dans une analyse en composantes principales: comparaison avec le modele du baton brise. J. Exp. Mar. Biol. Ecol. 25:67–75.

Fry, D.H. 1961. King salmon spawning stocks of the California Central Valley, 1940– 1959. California Fish and Game 47(1):55–71.

Ginetz, R.M., and P.A. Larkin. 1976. Factors affecting rainbow trout (*Salmo gairdneri*) predation on migrant fry of sockeye salmon (*Oncorhynchus nerka*). J. Fish. Res. Board Can. 33(1):19–24.

Gotelli, N.J and A.M. Ellison. 2004. A Primer of Ecological Statistics. Sinauer Associates, Inc. Publishers, Sunderland, Massachusetts. pp. 406–414.

Gregory, R.S. 1993. Effect of Turbidity on the Predator Avoidance Behaviour of Juvenile Chinook Salmon (*Oncorhynchus tshawytscha*). Can. J. Fish. Aquat. Sci. 50(2):241–246.

Gregory, R.S. and C.D. Levings. 1998. Turbidity Reduces Predation on Migrating Juvenile Pacific Salmon. Trans. Am. Fish. Soc. 127(2):275–285.

Grossman, G.D., T. Essington, B. Johnson, J. Miller, N.E. Monsen, and T.N. Pearsons. 2013. Effects of fish predation on salmonids in the Sacramento River–San Joaquin Delta and associated ecosystems. Panel final report 71 p. Sacramento (CA): CDFW, Delta Stewardship Council, and National Marine Fisheries Service. Available at: http://www.dfg.ca.gov/erp/predation.asp

Hall, R.O., J.B. Wallace, and S.L. Eggert. 2000. Organic matter flow in stream food webs with reduced detrital resource base. Ecology, 81(12):3445–3463.

Helena, B., R. Pardo, M. Vega, E. Barrado, J.M. Fernandez, and L. Fernandez. 2000. Temporal evolution of groundwater composition in an alluvial aquifer (Pisuerga river, Spain) by principal component analysis. Water Res. 34(3).

Helfield, J.M. and R.J. Naiman. 2001. Effects of Salmon-Derived Nitrogen on Riparian Forest Growth and Implications for Stream Productivity. Ecology, 82(9):2403–2409.

Henderson, M.J, I.S. Iglesias, C.J. Michel, A.J. Ammann, and D.D. Huff. 2019. Estimating spatial–temporal differences in Chinook salmon outmigration survival with habitat and predation-related covariates. Can. J. Fish. Aquat. Sci. 76(9):1549–1561.

Hennig, C. 2020. fpc: Flexible Procedures for Clustering. R package version 2.2-5. Available at https://CRAN.R-project.org/package=fpc

Higgins, J.V., M.T. Bryer, M.L. Khoury, and T.W. Fitzhugh. 2005. A Freshwater Classification Approach for Biodiversity Conservation Planning. Conserv. Biol. 19(2):432–445.

Jackson, M.C., O.L.F. Weyl, F. Altermatt, I. Durance, N. Friberg, A.J. Dumbrell, J.J. Piggott, S.D. Tiegs, K. Tockner, C.B. Krug, P.W. Leadley, and G. Woodward. 2016. Recommendations for the Next Generation of Global Freshwater Biological Monitoring Tools. *In* Advances in Ecological Research. *Edited by* A. J. Dumbrell, R. L. Kordas, and G. Woodward. Academic Press. **55**: 615–636.

Jeffres, C.A., Holmes, E.J., Sommer, T.R., and Katz, J.V.E. 2020. Detrital food web contributes to aquatic ecosystem productivity and rapid salmon growth in a managed floodplain. PLoS ONE 15(9): e0216019. https://doi.org/10.1371/journal.pone.0216019

Katz, J., P.B. Moyle, R.M. Quiñones, J. Israel, and S. Purdy. 2013. Impending extinction of salmon, steelhead, and trout (Salmonidae) in California. Environ. Biol. Fishes. 96(10):1169– 1186.

Lady, J.M. and J.R. Skalski. 2009. USER 4: User-Specified Estimation Routine. School of Aquatic and Fisheries Sciences, University of Washington, Seattle. Available at: http://www.cbr.washington.edu/analysis/apps/user/.

Lucas, L.V., J.E. Cloern, J.K. Thompson, and N.E. Monsen. 2002. Functional Variability of Habitats Within the Sacramento–San Joaquin Delta: Restoration Implications. Ecol. Appl. 12(5):1528–1547.

Lund, J., E. Hank, W. Fleenor, R. Howitt, J. Mount, and P.B. Moyle. 2007. Envisioning Futures for the Sacramento-San Joaquin Delta. Public Policy Institute of California: San Francisco.

Lyons, J., T. Zorn, J. Stewart, P. Seelbach, K. Wehrly, and L. Wang. 2009. Defining and Characterizing Coolwater Streams and Their Fish Assemblages in Michigan and Wisconsin, USA, North American Journal of Fisheries Management, 29:4, 1130–1151, doi: 10.1577/M08-118.1.

Maechler, M., P. Rousseeuw, A. Struyf, M. Hubert, and K. Hornik. 2019. *cluster*: Cluster Analysis Basics and Extensions. R package version 2.1.0.

Magnuson, J.J., L.B. Crowder, and P.A. Medvick. 1979. Temperature as an Ecological Resource. Am. Zool. 19(1):331–343.

McCullough, D., S. Spalding, D. Sturdevant, and M. Hicks. 2001. Summary of Technical Literature Examining the Physiological Effects of Temperature on Salmonids. EPA Issue Paper 5, EPA-910-D-01-005.

Michel, C.J., A.J. Ammann, S.T. Lindley, P.T. Sandstrom, E.D. Chapman, M.J. Thomas, G. P. Singer, A.P. Klimley, and R.B. MacFarlane. 2015. Chinook salmon outmigration survival in wet and dry years in California’s Sacramento River. Can. J. Fish. Aquat. Sci. 72(11):1749–1759.

Michel, C.J., C.M. Loomis, M.J. Henderson, J.M. Smith, N.J. Demetras, I.S. Iglesias, B.M. Lehman, and D.D. Huff. 2019. Linking predation mortality to predator density and survival for outmigrating Chinook Salmon and Steelhead in the lower San Joaquin River and South Delta. SWFSC for CDFW. E1696020, 58 p.

Michel, C.J. 2019. Decoupling outmigration from marine survival indicates outsized influence of streamflow on cohort success for California’s Chinook salmon populations. Can. J. Fish. Aquat. Sci. 76(8): 1398–1410. doi:10.1139/cjfas-2018-0140.

Moyle, P.B. 2002. Inland Fishes of California. UC Press. Berkeley. 502 pp

Moyle, P.B., R. Lusardi, P. Samuel, and J. Katz. 2017. State of the Salmonids: Status of California’s Emblematic Fishes 2017. Center for Watershed Sciences, University of California, Davis and California Trout, San Francisco, CA. 579 pp.

Natural Resource Defense Council v. Kirk Rodgers, as Regional Director of the United States Bureau of Reclamation, CIV-88-1658 LKK/GGH. 2006. Available at http://52.53.144.83/?wpfb_dl=9

Nislow, K.H. and J.D Armstrong. 2012. Towards a life-history-based management framework for the effects of flow on juvenile salmonids in streams and rivers. Fisheries Management Ecology, 9:451–463.

Pahl-Wostl, C. 2007. Transitions towards adaptive management of water facing climate and global change. Water Resour. Manag. 21(1):49–62.

Pedersen, E.J., Miller, D.L., Simpson, G.L., and Ross, N. 2019. Hierarchical generalized additive models in ecology: An introduction with mgcv. PeerJ, 7, e6876.

Peres-Neto, P.R., D.A. Jackson, and K.M. Somers. 2003. Giving Meaningful Interpretation to Ordination Axes: Assessing Loading Significance in Principal Component Analysis. Ecology, 84(9):2347–2363.

Perry, R.W., J.R. Skalski, P.L. Brandes, P.T. Sandstrom, A.P. Klimley, A. Ammann, and B. MacFarlane. 2010. Estimating Survival and Migration Route Probabilities of Juvenile Chinook Salmon in the Sacramento–San Joaquin River Delta. N. Am. J. Fish. Manage. 30(1):142–156.

Perry, R.W., A.C. Pope, J.G. Romine, P.L. Brandes, J.R. Burau, A.R. Blake, A.J. Ammann, and C.J. Michel. 2018. Flow-mediated effects on travel time, routing, and survival of juvenile Chinook salmon in a spatially complex, tidally forced river delta. Can. J. Fish. Aquat. Sci. 75(1): 1886–1901.

Petersen, J.H. and J.F. Kitchell. 2001. Climate regimes and water temperature changes in the Columbia River: bioenergetic implications for predators of juvenile salmon. Can. J. Fish. Aquat. Sci. 58:1831–1841.

Portz, D.E., S. Root, and C. Hueth. 2013. Central Valley Steelhead Monitoring Plan for the San Joaquin River Restoration Area: 2013 Monitoring Results for National Marine Fisheries Service Permit 16608. Technical Report. Available at http://www.restoresjr.net/?wpfb_dl=674\

R Core Team. 2017. R: A language and environment for statistical computing. R Foundation for Statistical Computing, Vienna, Austria. URL https://www.R-project.org/.

Robinson, A., A. Richey, J.E. Cloern, K.E. Boyer, J. Burau, E. Canuel, J. DeGeorge, J.Z. Drexler, L. Grenier, R.L. Grossinger, E.R. Howe, R. Kneib, R.J. Naiman, A. Mueller-Solger, J.L. Pinckney, D. Schoellhamer, and C. Simenstad. 2016. Primary Production in the Sacramento-San Joaquin Delta: A science strategy to quantify change and identify future potential. SFEI Contribution No. 781.

Rypel, A.L. 2014. The Cold-Water Connection: Bergmann’s Rule in North American Freshwater Fishes. The American Naturalist, 183(1):147–156.

Rypel, A.L., T.D. Simonson, D.L. Oele, J.D.T. Griffin, T.P. Parks, D. Seibel, C.M. Roberts, S. Toshner, L.S. Tate, and J. Lyons. 2019. Flexible Classification of Wisconsin Lakes for Improved Fisheries Conservation and Management. Fisheries, 44(5):225–238.

Safavian, S.R. and D. Landgrebe. 1991. A survey of decision tree classifier methodology. IEEE Transactions on Systems, Man, and Cybernetics, 21(3):660–674.

Sandstrom, P.T., D.L. Smith, and B. Mulvey. 2013. Two-dimensional (2-D) acoustic fish tracking at River Mile 85, Sacramento River, California. US Army Corps of Engineers, Engineer Research and Development Center.

Sass, G.G., A.L. Rypel, and J. Stafford. 2017. Inland Fisheries Habitat Management: Lessons Learned from Wildlife Ecology and a Proposal for Change. Fisheries, 42:197–209.

Schreck, C.B., M.F. Solazzi, S.L. Johnson, and T.E. Nickelson, 1989. Transportation stress affects performance of Coho Salmon, *Oncorhynchus kisutch*. Aquaculture, 82(1):15–20. https://doi.org/10.1016/0044-8486(89)90391-8.

Schupp, D.H. 1992. An Ecological Classification of Minnesota Lakes with Associated Fish Communities. Minnesota DNR Investigational Report (172).

Singer, G.P., A.R. Hearn, E.D. Chapman, M.L. Peterson, P.E. LaCivita, W.N. Brostoff, A. Bremner, and A.P. Klimley. 2013. Interannual variation of reach specific migratory success for Sacramento River hatchery yearling late-fall run Chinook salmon (*Oncorhynchus tshawytscha*) and steelhead trout (*Oncorhynchus mykiss*). Environ. Biol. Fishes. 96(2–3):363–379.

SJRRP (San Joaquin River Restoration Program). 2018. San Joaquin River Restoration Program 2017-2018 Annual Report. Technical Document. Available at http://www.restoresjr.net/science/reports/.

SJRRP (San Joaquin River Restoration Program). 2019. Vegetation Monitoring along the San Joaquin River. Technical Document. Available at http://www.restoresjr.net/science/fisheries-and-habitat/

Sommer, T., C. Armor, R. Baxter, R. Breuer, L. Brown, M. Chotkowski, S. Culberson, F. Feyrer, M. Gingras, B. Herbold, W. Kimmerer, A. Mueller-Solger, M. Nobriga, and K. Souza. 2007. The Collapse of Pelagic Fishes in the Upper San Francisco Estuary. Fisheries, 32(6):270–277.

Steel, E.A., D.W. Jensen, K.M. Burnett, K. Christiansen, J.C. Firman, B.E. Feist, K.J. Anlauf, and D.P. Larsen. 2012. Landscape characteristics and Coho Salmon (*Oncorhynchus kisutch*) distributions: explaining abundance versus occupancy. Can. J. Fish. Aquat. Sci. 69(3):457–468.

Thorp, J.H., M.C. Thomas, and M.D. Delong. 2008. The Riverine Ecosystem Synthesis: Toward Conceptual Cohesiveness in River Science. Academic Press.

Tyers, M. 2020. *Riverdist*: River Network Distance Computation and Applications. R package version 0.15.2.

Van den Brink, P.J., C.B. Choung, W. Landis, M. Mayer-Pinto, V. Pettigrove, P. Scanes, R. Smith, and J. Stauber. 2016. New approaches to the ecological risk assessment of multiple stressors. Mar. Freshw. Res. 67(4):429.

Vogel, David A. 2010. Evaluation of Acoustic-Tagged Juvenile Chinook Salmon Movements in the Sacramento-San Joaquin Delta during the 2009 Vernalis Adaptive Management Program. Technical Report to the California Water Resources Control Board. Available at: www.sjrg.org/technicalreport/.

Warren, C.E., P. Doudoroff, and D.L. Shumway. 1973. Development of Dissolved Oxygen Criteria for Freshwater Fish. Western Fish Toxicology Laboratory for the EPA Office of Research and Monitoring. EPA-R3-73-019.

Wehrly, K.E., J.E. Breck, L. Wang, and L. Szabo-Kraft. 2012. A Landscape-Based Classification of Fish Assemblages in Sampled and Unsampled Lakes. Trans. Am. Fish. Soc. 141(2):414–425.

Weitkamp, L.A. 1994. A Review of the effects of dams on the Columbia River Estuarine Environment, with special reference to Salmonids. Bonneville Power Administration.

Wiens, J.A. 2002. Riverine landscapes: taking landscape ecology into the water. Freshw. Biol. 47(4):501–515.

Wohl, E., S.N. Lane, and A.C. Wilcox. 2015. The science and practice of river restoration. Water Resources Research, 51:5974–5997.

Wood, S.N. 2011. Fast stable restricted maximum likelihood and marginal likelihood estimation of semiparametric generalized linear models. Journal of the Royal Statistical Society (B), 73(1):3–36.

Yohannes, Y. and P. Webb. 1999. Classification and Regression Trees, CART: A User Manual for Identifying Indicators of Vulnerability to Famine and Chronic Food Insecurity. International Food Policy Research Institute: Washington, D.C.

Yoshiyama, R.M., F.W. Fisher, and P.B. Moyle. 1998. Historical Abundance and Decline of Chinook Salmon in the Central Valley Region of California. N. Am. J. Fish. Manage 18(3): 487–521.

Yoshiyama, R.M., E.R. Gerstung, F.W. Fisher, and P.B. Moyle. 2000. Chinook Salmon in the California Central Valley: An Assessment. Fisheries Management, 25(2):15.

Yoshiyama, R.M, E.R. Gerstung, F.W. Fisher, and P.B. Moyle. 2001. Historical and Present Distribution of Chinook Salmon in the Central Valley Drainage of California. California Department of Fish and Game Fish Bulletin. 179(1).

Zabel, R.W. and S. Achord. 2004. Relating size of juveniles to survival within and among populations of chinook salmon. Ecology, 85(3):795–806.

Zeug, S.C., J. Wiesenfeld, K. Sellheim, A. Brodsky, and J.E. Merz. 2019. Assessment of Juvenile Chinook Salmon Rearing Habitat Potential Prior to Species Reintroduction. N. Am. J. Fish. Manage. 39(4):762–777.

Zillig, K.W., Lusardi, R.A., Moyle, P.B., and N.A. Fangue. 2021. One size does not fit all: variation in thermal eco-physiology among Pacific salmonids. Rev Fish Biol Fisheries 31: 95– 114. https://doi.org/10.1007/s11160-020-09632-w

