## Supplementary Materials for: Survival of a threatened salmon is linked to spatial variability in river conditions for "Survival of a threatened salmon is linked to spatial variability in river conditions"

Colby L. Hause^1^, Gabriel P. Singer^1^, Rebecca A. Buchanan^2^, Dennis E. Cocherell^3^, Nann A. Fangue^3^, and Andrew L. Rypel^3, 4^

1: California Department of Fish and Wildlife, 1010 Riverside Parkway, West Sacramento, CA 95605.

2: School of Aquatic and Fishery Sciences, University of Washington, 1325 Fourth Avenue, Suite 1515, Seattle, Washington 98101-2540.

3: Department of Wildlife, Fish, and Conservation Biology, One Shields Avenue, University of California, Davis, CA 95616.

4: Center for Watershed Sciences, University of California, Davis, CA 95616.

**Supplemental Methods**

**FLAMe Validation**

To validate Suna and YSI Exo2 measurements of NO_3_, chlorophyll-$\alpha$ and DOC, we collected discrete water samples at 12 sites along the FLAMe transect for lab analysis. Once on site, 1000 mL brown Nalgene bottles were rinsed, filled with surface water, and placed on ice. All samples were processed in the lab within 12 hours of collection. For chlorophyll-α, river water was filtered onto 1.2 µm VWR glass fiber filters and frozen until analysis. Samples were analyzed using the 90% acetone method via a spectrophotometer at absorbance wavelengths of 750, 665, and 664. To validate NO_3_ measurements, river water was filtered (1.2 µm) into 20 mL plastic scintillation vials and immediately frozen after processing. Lab NO_3_ samples were analyzed by the hydrazine method, which uses a hydrazine-copper solution to reduce nitrate (NO_3_) to nitrite (NO_2_) followed by color development using a diazotization-coupling reaction. For DOC analysis, whole water samples were poured into 100 mL amber glass vials treated with phosphoric acid preservative. Samples were stored on ice and sent to Alpha Analytical (Sacramento, CA) for analysis using Standard Method 5310C, which is a persulfate-ultraviolet oxidation method. In situ measurements of variables that could not be lab-analyzed (DO, pH, temperature, turbidity and specific conductivity) were recorded on a YSI 6820 at each station to validate YSI Exo2 measurements. The YSI Exo2 and YSI 6820 were calibrated prior to each sampling event and functionality of the Suna sensor was confirmed using deionized water measurements.

To further validate both the filtering procedure and the lab analysis results, all lab samples included one duplicate and one field blank for each sampling event (n = 3). Duplicate samples should yield similar results, and large differences between duplicate sample readings indicate that an error during lab analysis or sample collection could have occurred. Field blanks are samples filtered using deionized (DI) water only, and results that deviate from zero indicate potential sample contamination during the filtering process or errors during lab analysis. We assessed the relationship between validated (via lab analysis or *in situ* YSI 6820 readings) and FLAMe-collected measurements using linear regression analysis (R Core Team 2017). Highly correlated relationships (r^2^ value close to 1) indicated accurate characterization of that parameter by the FLAMe sensor.

**Fish tagging and releases**

A total of 750 spring-run Chinook salmon smolts were obtained from SCARF (Salmon Conservation and Research Facility) and surgically implanted with JSATS acoustic transmitters (ATS model SS400, 216 mg, 3.38mm x 15.0 mm). Food was withheld from all fish at least three days prior to tagging, and not provided again until at least 24 hours after tagging. Smolts ranging in size from 73-93 mm (mean fork length (mm) = 81.30 ± 4.02 SD, mean weight (g) = 5.79 ± 0.89 SD) were anesthetized with a buffered solution of tricaine methanesulfonate (4.32 g sodium bicarbonate, 1.02 g MS-222) at a concentration of 90 mg L^-1^ in accordance with UC Davis *Institutional Animal Care and Use Committee (IACUC) Protocol Number 21614*. Once fish lost the ability to maintain equilibrium (stage 4 anesthesia; Summerfelt and Smith 1990), they were weighed (grams), measured (fork length, mm), and visually inspected for any abnormalities. Fish were then placed ventral side up onto a foam surgery cradle and kept moist with a maintenance dose (i.e. surgery bath) of anesthesia (concentration 30 mg L^-1^, Singer et al. in prep^[1]^). Anesthetic and surgery bath solutions were periodically checked to ensure adequate oxygen saturation and that temperatures remained ±2˚C of the ambient water temperature. Acoustic tags were soaked in a bath of nolvasan dilution for disinfection purposes and rinsed with deionized water. A 5-8 mm incision was made parallel to the mid-ventral line and ended 3-5 mm anterior to the pelvic girdle. Tags were inserted into the coelom of the fish and secured with one suture (PDS 5/0 monofilament) tied with a 2x2 surgeon’s knot. All fish weighed at least 4.2 grams to maintain a tag burden of 5% or less. Upon completing surgeries, fish recovered in a bucket of water and were then returned to a holding tank. A total of 350 fish were tagged for each of two release groups and another 50 fish were tagged as part of a concurrent tag-effects study. The 50 tag-effects fish were placed in a separate holding tank to monitor tag battery life, shedding rates, and any surgery-related mortalities over the course of the study window (February- mid June).

After a minimum 24-hour holding period post-surgery, 350 tagged smolts were transported in a CDFW-operated hauling tank along with 80,000-100,000 untagged smolts for release at the upper release site in the SJRRP Restoration Area (Fig.1). During transport, temperature and oxygen levels were checked every 30-60 minutes and oxygen was adjusted to maintain ~100% saturation. Upon arrival to the upper release location, fish were acclimated to river water temperatures by releasing water from the hauling tank while pumping river water in at a rate of 1˚C increase per hour until hauling tank temperatures were ±2˚C of ambient river water. Acclimation procedures lasted 3 hours to account for the 3°C difference in river water temperature from the hauling tank, after which fish were released at sundown.

Movements of tagged fish were tracked using real-time receiver technology located at Hills Ferry (A4, Fig.1) to coordinate timing of the Delta release downriver. The purpose of the Delta release was to ensure our ability to estimate survival to the ocean in the event of low survival from the upstream release group, so it was important that fish from each group enter the Delta around the same time. Once detections at Hills Ferry peaked, the second group of 350 fish were transported from the holding tank at SIRF (Satellite Incubation and Rearing Facility) to the Delta release site located at Durham Ferry (A7, Fig. 1) on March 12, 2019. These fish were transported in a smaller hauling tank along with ~500 untagged smolts. Transport and acclimation protocols remained consistent with those of the upper release, and fish were acclimated for 1 hour before they were released into the river after sundown.

**FLAMe Data Processing**

Raw water quality data from FLAMe surveys were post-processed to remove erroneous readings and blank records. Erroneous readings were quantitatively identified and removed using the median absolute deviation (MAD) and median calculated within a rolling window of 20-100 observations, depending on the variable. Maximum allowable MAD values away from the median within each window were set as threshold criteria (ranging from 3-7 MADs) for each environmental variable. Data that fell outside these ranges were flagged and manually removed upon visual inspection. NAs (blank readings) were recorded when the signal output from sensors was slower than the collection timing of the datalogger (1 collection per second) or when a sensor failed to take a measurement during a collection. Values that resulted from sampling errors were also manually removed from the dataset, which included periods of time when mud or debris clogged the flow-through system or when the intake pipe introduced air into the sensor chambers. Cluster analysis requires that every sample contains values for all variables measured (Ryberg 2006), and therefore any coordinate with missing water quality information was removed from the impacted transect. GPS coordinates were converted to river kilometers values using the R package *nhdplusTools* (v0.3.13; Blodgett 2018) and all data points were snapped to the centerline of the river using the R package *riverdist* (v0.15.0; Tyers 2020).

**Acoustic Telemetry Data Processing**

Previous studies suggest predation on acoustic-tagged salmon smolts can be high, which has potential to bias survival results if unaccounted for (Vogel 2010; Buchanan et al. 2013). We applied a behavior-based predator filter that evaluated movement patterns for each tag according to our assumptions of differences between predator and smolt movements. Examples of patterns assumed to be indicative of predators include upstream movement against localized flow (either river or tidal flow) and movement in-and-out of receiver range but within the same general area for long periods of time (Vogel 2010; Buchanan et al. 2013). Our filtering process started with the application of a broad predator filter to flag upstream movements greater than 16 km, starting at the last downstream detection location. This large threshold was selected to prevent tidally-induced movements from being truncated within the interior Delta, which was typically < 16 km based on previous observations (Singer et al. in prep^1^).

Next, detections for each tag were plotted and visually assessed for irregular movement patterns and manually cleaned. Tags that recorded upstream movement over shorter distances were analyzed based on location, discharge, and tidal influence. If the detection sequence was within an area of tidal influence and followed patterns of upstream movement on the flood tide and downstream movement on the ebb (Perry et al. 2010), this movement was considered consistent with smolt behavior and only the furthest downstream detections were retained. On many occasions, back and forth movement was detected in the interior Delta between water operation facilities (D1, E1, E2) and Old River and Middle River sites at Highway 4 (B2) (Fig. 1). In most cases, these movements were considered to be indicative of predator consumption (Vogel 2010; Buchanan et al. 2013), and truncated at the last location the salmon was assumed to be alive. The resulting detection histories for each tag represented the chronological order of receiver locations and final downstream-directed detections during outmigration.

**Supplemental Results**

**FLAMe Sample Validation**

Water quality variables generally exhibited strong linear relationships between FLAMe and in situ/lab validation measurements (*r^2^* range = 0.599-0.999), with the exception of pH and chlorophyll-$\alpha$ (*r^2^* range = 0.068-0.245) (Fig. S2). We observed a weak linear relationship (*r^2^* = 0.068, Fig. S2) between FLAMe YSI Exo and YSI 6820 pH values, and a relatively high difference between sensor readings (± 0.48) in relation to the range of the two probes (7.16 – 7.81). There are a couple of possible explanations for this result. The YSI 6820 contained an older pH sensor model, which may have decreased sensor accuracy. It is likely that the older model also had a longer sensor response time than the newer FLAMe pH probe, thus providing an additional source of variation between the probes. However, due to the overall variable nature of pH sensors, a 0.48 difference between sensor measurements did not cause alarm. Both probes characterized the river within the same pH unit, and across a range that we might expect to observe in the natural environment. Future studies should better account for probe measurement error by (a) performing validation measurements with similar sensor models and/or (b) conducting sensor response time experiments to better account for variation in acclimation time between different sensor models.

There was also weak correlation (*r^2^* = 0.245) between the FLAMe and lab analysis readings for chlorophyll-$\alpha$ concentrations (Fig. S2). We do not think that these results reflect inaccurate chlorophyll-$\alpha$ readings on the FLAMe or improper calibration of the probe, but rather an issue during lab analysis. Duplicate sample values for chlorophyll-$\alpha$ resulted in large differences (ranging 0.2-11.21 mg m^3 -1^) across all sampling events, and field blanks (which should result in a chlorophyll-$\alpha$ value of 0) were consistently high (3.2 – 14.42 mg m^3 -1^). Chlorophyll-$\alpha$ values measured by the FLAMe were similar to CDEC measurements at stations equipped with chlorophyll-$\alpha$ sensors, thus providing further support for accurate measurements by the FLAMe chlorophyll-$\alpha$ probe.

**Tag Effects**

Battery life of acoustic transmitters, tag shedding, and fish mortality were monitored in 50 tagged fish held at SIRF from the start of tagging until all receivers were removed from the river in mid-June. 98% of the batteries in the 50 tags were still operational by the end of the time period when fish were detected in the receiver array. Under the assumption of 100% tag survival, the estimated probability of survival ($\hat{S}$) to Golden Gate bridge (A17, Fig.1) was 0.050 (± 0.009 $\hat{SE}$). Under 98% tag survival, the adjusted estimate would have been $\hat{S}$ = 0.051, which is well within the standard error of the unadjusted estimate. While no fish mortality was observed, three tags were recorded to have been shed from fish held at the hatchery over the study period. This would have adjusted survival estimates to Golden Gate bridge from 0.050 to 0.053, also within the standard error of the unadjusted estimate. Therefore, no adjustments to survival estimates were calculated for premature tag failure or shedding.

**Supplemental Tables**

Table S1. Summary of water chemistry patterns through each cluster used in fitting model M_2_, as determined by AHC analysis on the mean values of the three water quality transects.

| **Cluster** | **Reaches** | **Region** | **Water Chemistry Description** |
| --- | --- | --- | --- |
| C1 | A1 - A4 | Restoration Area | Highest observed values of chlorophyll-a, turbidity, fDOM, specific conductivity and temperature. Lowest DO levels and NO_3_ within mid-range. |
| C2 | A5 | Upper River | Higher variance in the distribution of NO_3_ (IQR 33-40 uM), but median value was comparable to C1. Chlorophyll-$\alpha$, turbidty, fDOM, specific conductivity and temperature decreased from C1 levels while DO increased slightly. |
| C3 | A6 | Mid-River | Relatively high variation observed in all variables with the exception of turbidity. NO_3_ reached maximum values, but accompanied by high variance (IQR 47.8-59 uM). Chlorophyll-$\alpha$, fDOM, specific conductivity and temperature maintained similar median values to those in C2, while median turbidity decreased slightly. DO increased from C2 levels. |
| C4 | A7-A10 | Lower River | NO_3_, specific conductivity, chlorophyll-$\alpha$, and fDOM decreased from C3 levels and temperature reached its lowest mean value. Compared to C3, median turbidity levels decreased slightly and DO increased. |
| C5 | A11-A12 | Southern Delta | Lowest levels of NO_3_, specific conductivity and fDOM and highest recorded DO (>100% saturation). Chlorophyll-$\alpha$ and turbidity decreased from C4 levels while temperature increased by ~ 2 $^{\circ}$C. |
| C6 | A13 | Mainstem Central Delta | NO_3_, fDOM and specific conductivity remained low, though increasing slightly from C5 levels. Turbidity and chlorophyll-$\alpha$ reached lowest values while DO remained high (>100% saturation). Temperature increased from C5 levels, nearly approaching median values observed in C1 (15.9 $^{\circ}$C). |

Table S2. Squared PCA loadings across 2 principal components from the mean (A) and CV (B) datasets in relation to broken-stick model expected values. Loading significance was determined following steps outlined in Peres-Neto et al. (2003). First, component values were ranked across all 8 principle components (not shown) for comparison against expected values. Component values whose ranked squared loadings were greater than the respective expected values were identified as significant (indicated in bold). Only components 1 and 2 were considered by the broken‐stick criterion, as informed by results from the Kaiser Guttman criterion.

|  | Component (A) | | Component (B) | |
| --- | --- | --- | --- | --- |
| Variable | 1 | 2 | 1 | 2 |
| Chl-$\alpha$ | 0.1748 | 0.0062 | 0.1271 | 0.1459 |
| Turbidity | 0.0572 | 0.3411 | 0.1302 | 0.1184 |
| NO_3_ | 0.1478 | 0.0003 | 0.0683 | 0.2001 |
| Sp.Cond. | 0.1741 | 0.0139 | 0.1845 | 0.0459 |
| Temp | 0.0616 | **0.3704** | **0.2494** | 0.0085 |
| pH | 0.0572 | 0.2082 | 0.0683 | 0.2001 |
| fDOM | 0.1729 | 0.0104 | 0.0042 | **0.2316** |
| DO | 0.1543 | 0.0496 | 0.1619 | 0.0803 |
| *Expected* | **0.3397** | 0.2147 | 0.3397 | **0.2147** |

Table S3. Parameter estimates for the spatial (M_2_) and the geographic (M_1_) models. Parameters include survival (presented as per-km survival $\sigma$ and survival probability *S*), route selection $\psi$, and transition probability $\phi$. Reach length (km) is provided for the corresponding parameter when applicable. Parameters correspond to the model schematic (Fig. S1). S.E. = estimated standard error of *S*, $\psi$, or $\phi$ estimate.

| Model | Region | Cluster | Parameter | Estimate | S.E. | Length (km) | $\hat{\sigma}$ |
| --- | --- | --- | --- | --- | --- | --- | --- |
| M_2_ | Restoration Area | 1 | sA_1*_sA_2*_sA_3*_sA_4_ | 0.6104 | 0.0259 | 14.0 | 0.9642 |
| M_2_ | Upper River | 2 | sA_5_ | 0.7613 | 0.0298 | 17.4 | 0.9844 |
| M_2_ | Mid-River | 3 | sA6 | 0.8936 | 0.0245 | 29.3 | 0.9962 |
| M_2_ | Lower River | 4 | sA_7*_sA_8*_sA_9*_sA_10_ | 0.9409 | 0.0205 | 51.3 | 0.9988 |
| M_2_ | Southern Delta | 5 | sA_11*_sA_12_ | 0.9603 | 0.0224 | 18.2 | 0.9978 |
| M_2_ | Central Delta | 6 | sA_13_ | 0.6300 | 0.0422 | 14.4 | 0.9684 |
| M_1_ | Central Delta | - | sA_14_ | 0.4582 | 0.0515 | 29.0 | 0.9734 |
| M_1_ | Estuary | - | sA_15_ | 0.7643 | 0.0621 | 22.0 | 0.9878 |
| M_1_ | Estuary | - | sA_16_ | 0.9535 | 0.0321 | 19.3 | 0.9975 |
| M_1_ | Estuary | - | sA_17_$\dagger$ | 1 | - | 50.9 | 1 |
| M_1_ | Southern Delta | - | $\psi$A_1_ | 0.5920 | 0.0427 | - | - |
| M_1_ | Central Delta | - | $\psi$A_2_ | 0.7953 | 0.0837 | - | - |
| M_1_ | Central Delta | - | $\psi$F | 0.2047 | 0.0837 | - | - |
| M_1_ | Central Delta | - | sF_1_ | 0.1177 | 0.1199 | 30.9 | 0.9331 |
| M_1_ | Southern Delta | - | $\psi$B | 0.4080 | 0.0427 | - | - |
| M_1_ | Interior Delta | - | $\phi$B_1_E_1_ | 0.5825 | 0.0696 | 24.3 | - |
| M_1_ | Interior Delta | - | $\phi$B_1_D_1_ | 0.1101 | 0.0424 | 22.6 | - |
| M_1_ | Interior Delta | - | $\phi$B_1_B_2_ | 0.2386 | 0.0578 | 32.3 | - |
| M_1_ | Interior Delta | - | sE_1_ | 0.7245 | 0.0823 | 3.2 | 0.9027 |
| M_1_ | Interior Delta | - | sE_2_ | 0.1795 | 0.0615 | $\ddagger$ | $\ddagger$ |
| M_1_ | Interior Delta | - | sD_1_$\dagger$ | 0 | - | $\ddagger$ | $\ddagger$ |
| M_1_ | Interior Delta | - | sB_1_$\dagger$ | 0 | - | 43.2 | 0 |

$\dagger$ Parameter was fixed in model based on observed data

$\ddagger$ Pathway consists of transport via trucking, therefore no reach length available

**Supplemental Figures**

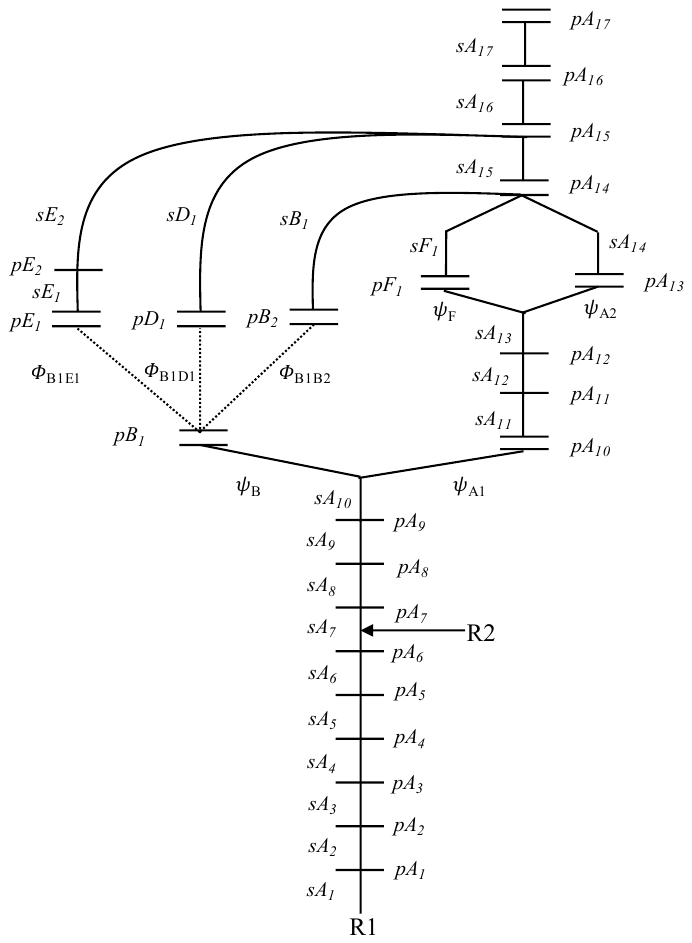

Figure S1. Schematic of the multistate mark-recapture model. Parameters include reach-specific survival (*S),* detection probability (*p*), route selection (𝜓), and transition probability ($\phi$). Horizontal lines represent acoustic receivers and parallel lines indicate dual receiver arrays. R1 and R2 mark locations of release 1 (upper release) and 2 (Delta release).

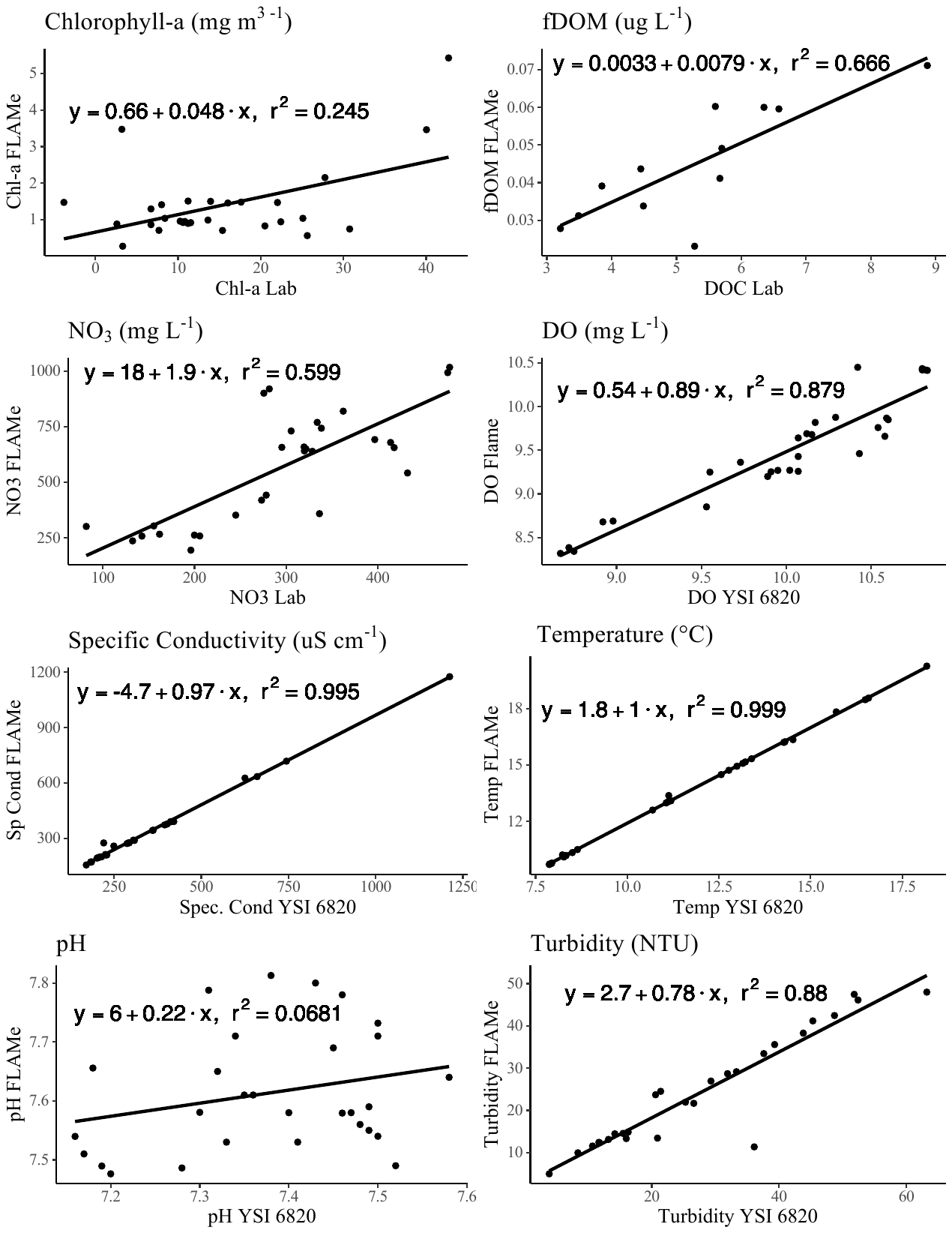

Figure S2. Relationship between FLAMe and validation measurements. Temperature, pH, DO, specific conductivity, and turbidity were validated via *in situ* measurements using a YSI 6820, while chlorophyll-$\alpha$, NO_3_, and fDOM variables were validated via lab analysis.

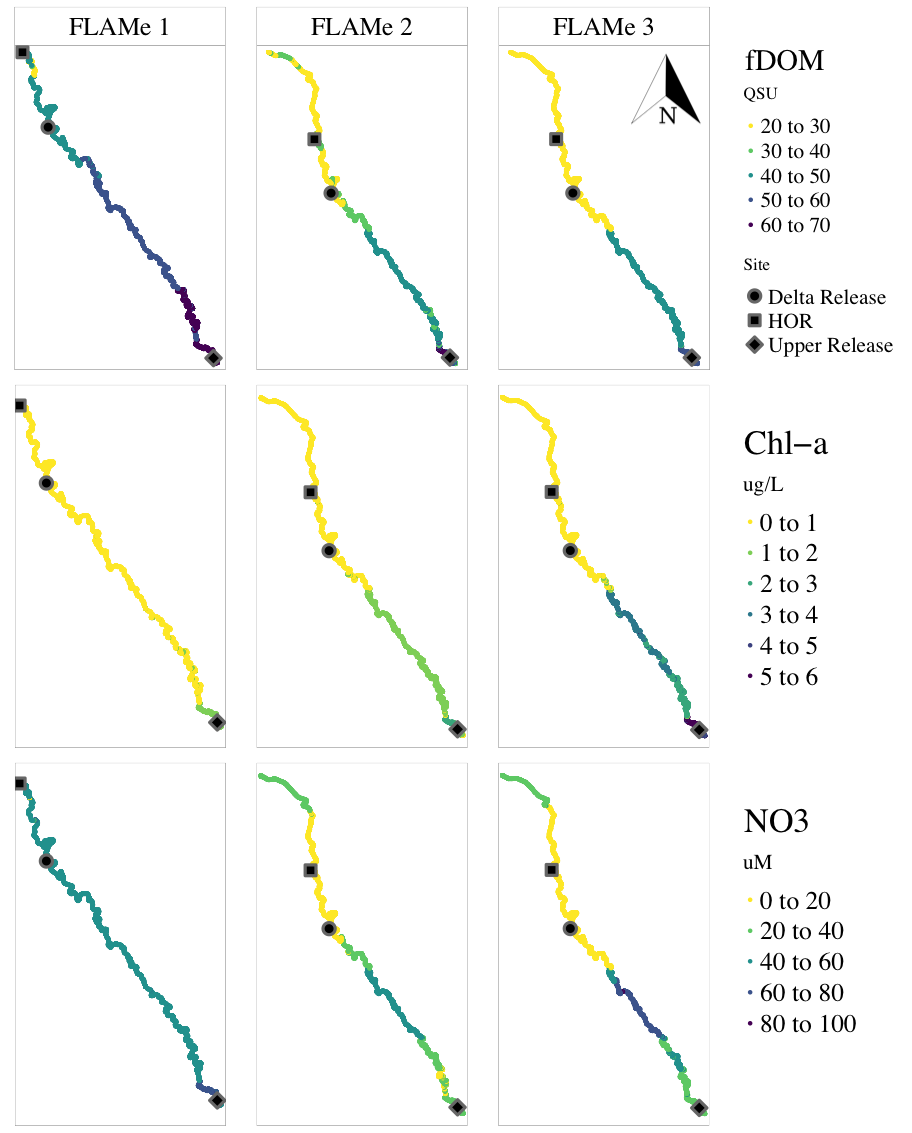

Figure S3a. Spatial distribution of fDOM, chlorophyll-$\alpha$, and NO_3_ values observed for each FLAMe transect (n= 3). See Table 2 for sampling dates. The three reference points on each map mark the locations of the upper release (diamond), Delta release (circle), and the head of Old River (HOR) junction (square).

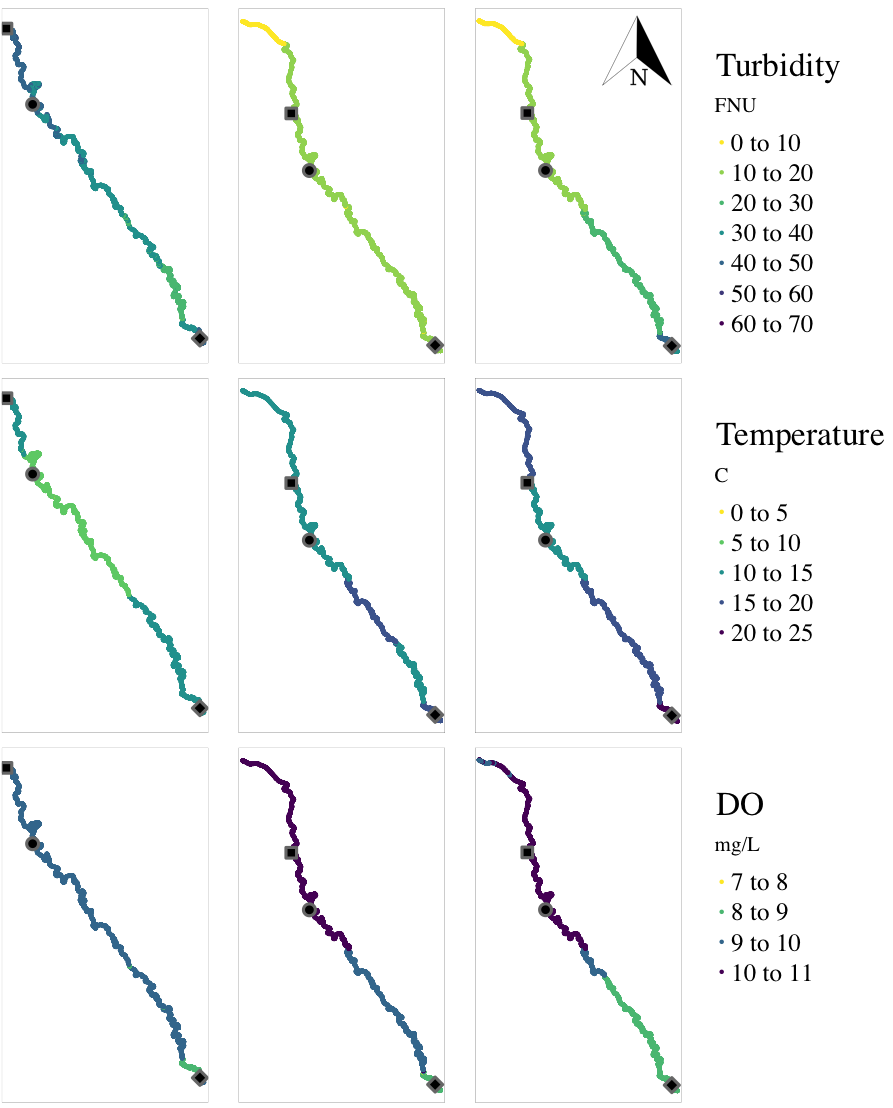
 Figure S3b. Spatial distribution of turbidity, temperature, and DO values observed for each FLAMe transect (n= 3). See Table 2 for sampling dates. The three reference points on each map mark the locations of the upper release (diamond), Delta release (circle), and the head of Old River (HOR) junction (square).

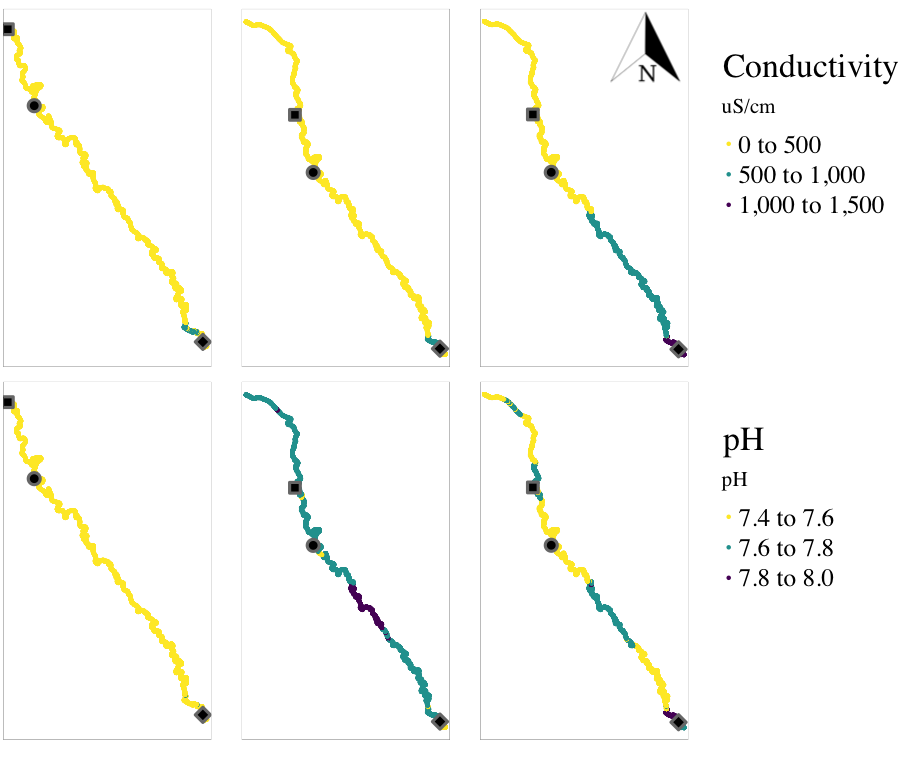

Figure S3c. Spatial distribution of specific conductivity and pH values observed for each FLAMe transect (n= 3). See Table 2 for sampling dates. The three reference points on each map mark the locations of the upper release (diamond), Delta release (circle), and the head of Old River (HOR) junction (square).

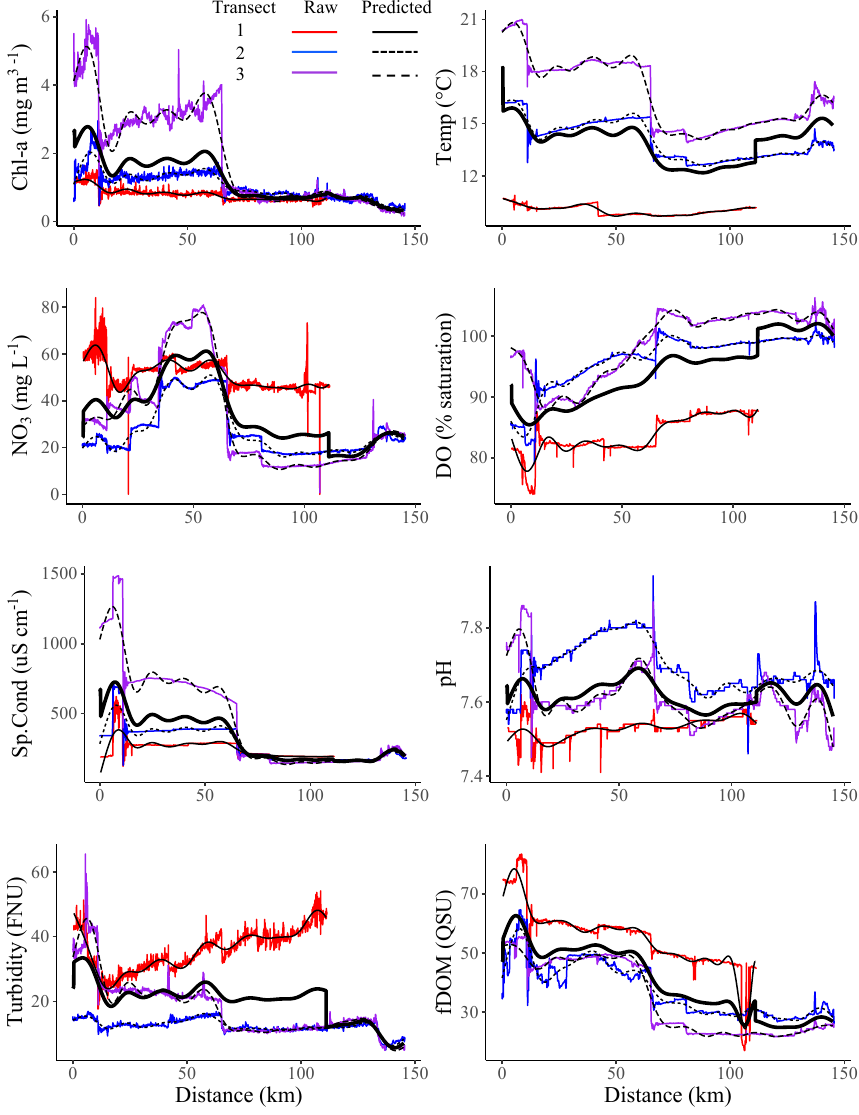

Figure S4. Raw vs predicted values from the GAMM analysis for each variable across the three transects. The bold line represents the mean value calculated from the predicted values, and the x axis represents distance from the upper release location (Fig. 1).

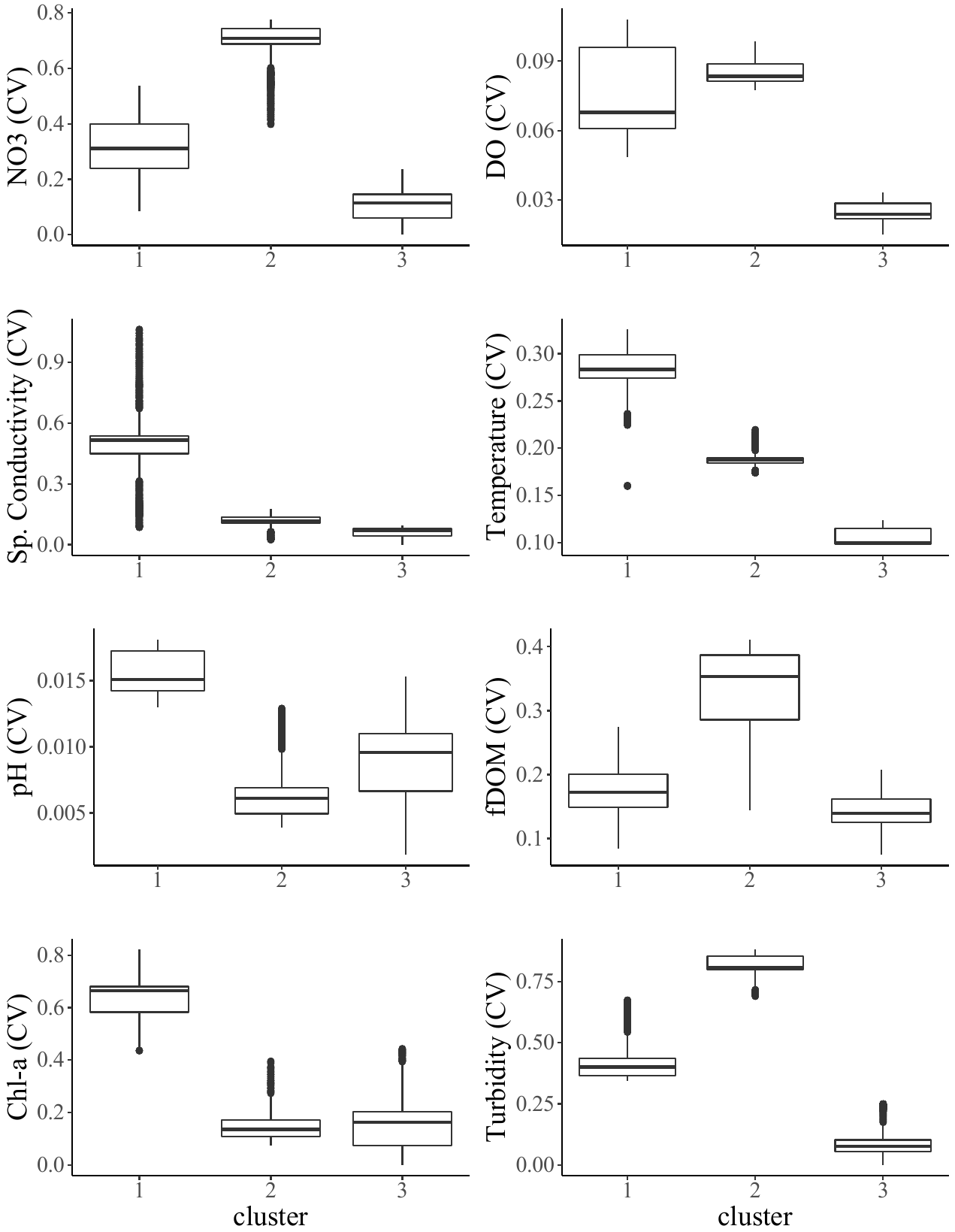

Figure S5. Coefficient of variation of water chemistry variables distributed across clusters, as determined by cluster analysis on the CV dataset. The bold horizontal lines represent median values, while the upper and lower edges of the boxes represent the 75^th^ (Q3) and 25^th^ (Q1) percentiles, respectively. The upper and lower ends of the vertical lines represent largest and smallest value no further than 1.5 *IQR (interquartile range, or Q3-Q1) from the 75^th^ and 25^th^ quantiles. Points beyond the end of the vertical lines represent outliers.

**Literature Cited**

Blodgett, D. 2018. *nhdplusTools:* Tools for Accessing and Working with the NHDPlus.<https://code.usgs.gov/water/nhdplusTools>.

Buchanan, R.A., J.R. Skalski, P.L. Brandes, and A. Fuller. 2013. Route Use and Survival of Juvenile Chinook Salmon through the San Joaquin River Delta. North American Journal of Fisheries Management **33**(1):216–229.

Perry, R.W., J.R. Skalski, P.L. Brandes, P.T. Sandstrom, A.P. Klimley, A. Ammann, and B. MacFarlane. 2010. Estimating Survival and Migration Route Probabilities of Juvenile Chinook Salmon in the Sacramento–San Joaquin River Delta. N. Am. J. Fish. Manage. **30**(1):142–156.

R Core Team. 2017. R: A Language and Environment for Statistical Computing, R Foundation for Statistical Computing, Vienna, Austria. Available at [http://www.R‐project.org/](http://www.r-project.org/).

Ryberg, K.R. 2006. Cluster Analysis of Water-Quality Data for Lake Sakakawea, Audubon Lake, and McClusky Canal, Central North Dakota, 1990-2003. Scientific Investigations Report.

Summerfelt, R. and L. Smith. 1990. Anesthesia, surgery, and related techniques. Pages 213–272 *in* C. B. Schreck and P. B. Moyle, editors. Methods for Fish Biology. American Fisheries Society, Bethesda, MD.

Tyers, M. 2020. *Riverdist*: River Network Distance Computation and Applications. R package version 0.15.2.

Vogel, David A. 2010. Evaluation of Acoustic-Tagged Juvenile Chinook Salmon Movements in the Sacramento-San Joaquin Delta during the 2009 Vernalis Adaptive Management Program. Technical Report to the California Water Resources Control Board. Available: [www.sjrg.org/technicalreport/](http://www.sjrg.org/technicalreport/).

1. Singer, G.P., C.L. Hause, E.D. Chapman, M.P. Pagel, A.P. Klimley, N.A. Fangue, and A.L.

   Rypel. Dynamics of juvenile survival for reintroduced spring-run Chinook Salmon smolts. *Manuscript in Preparation.*
